## Supplemental Figures for "Chromoanagenesis landscape in 10,000 TCGA patients"

**Figure S1. CNV frequency in the PCAWG-TCGA cohort**

**(a)** Box plot of number of deleted genes for each chromoanagenesis subtype. **(b)** Box plot of number of amplified genes for each chromoanagenesis subtype.

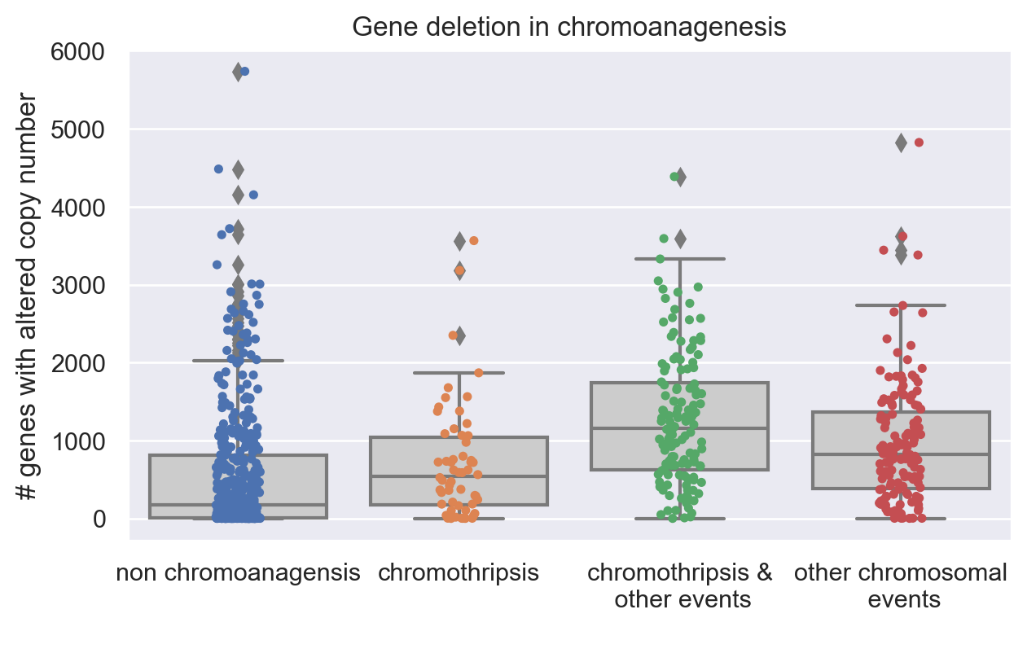
**a**

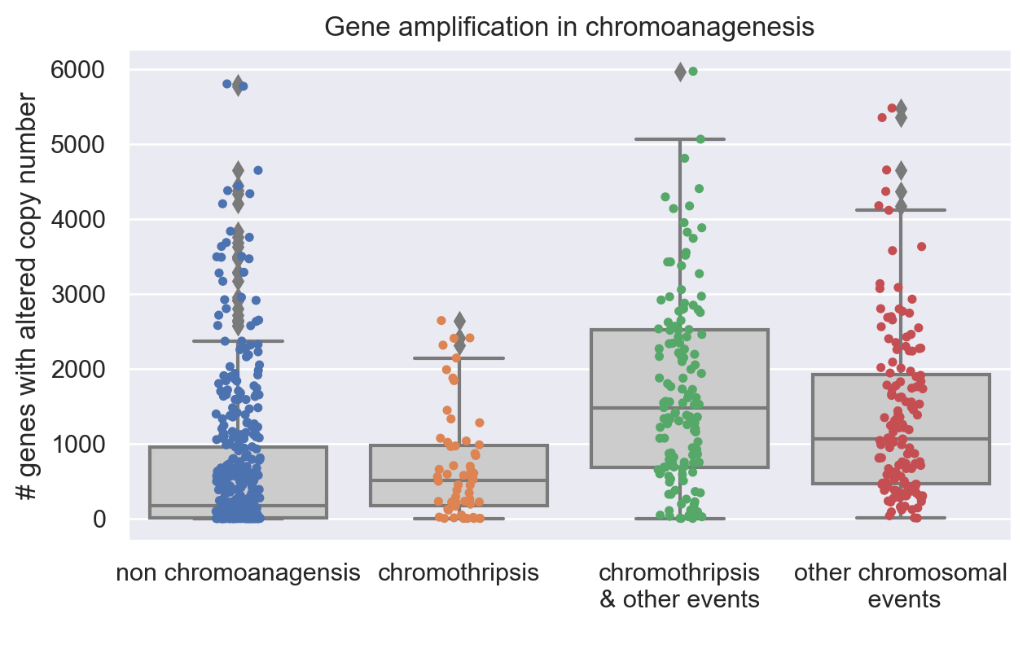
**b**

**Figure S2. BLCA Manhattan plots**

Genic Manhattan plot over Fisher’s exact test p-values between BLCA chromoanagenesis samples and non-chromoanagenesis samples. **(a)** Manhattan CNA (deletion or amplification) plot. **(b)** Manhattan Deletion plot. **(c)** Manhattan Amplification plot.

**a**
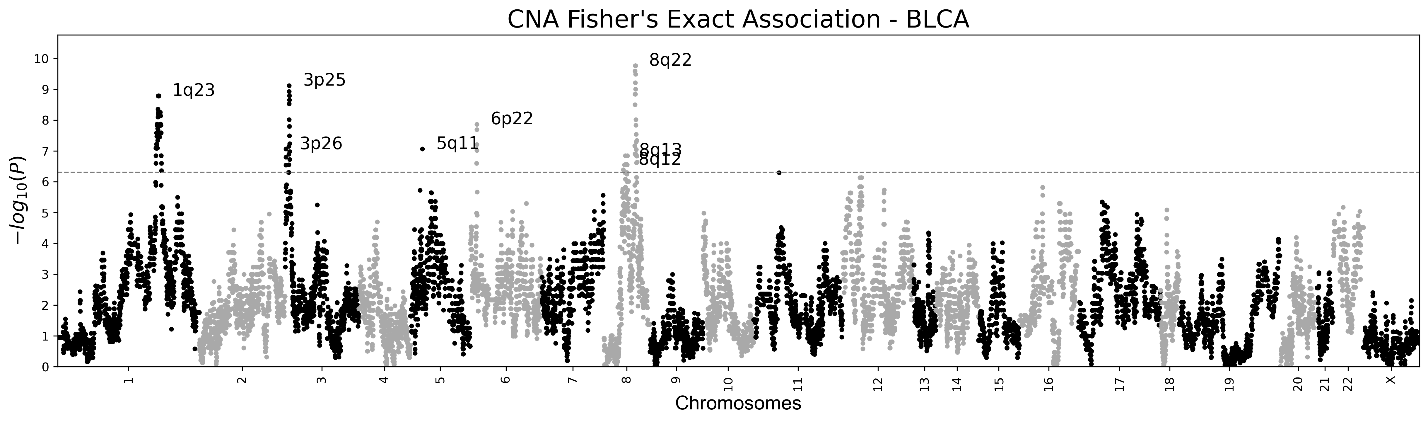

**b**
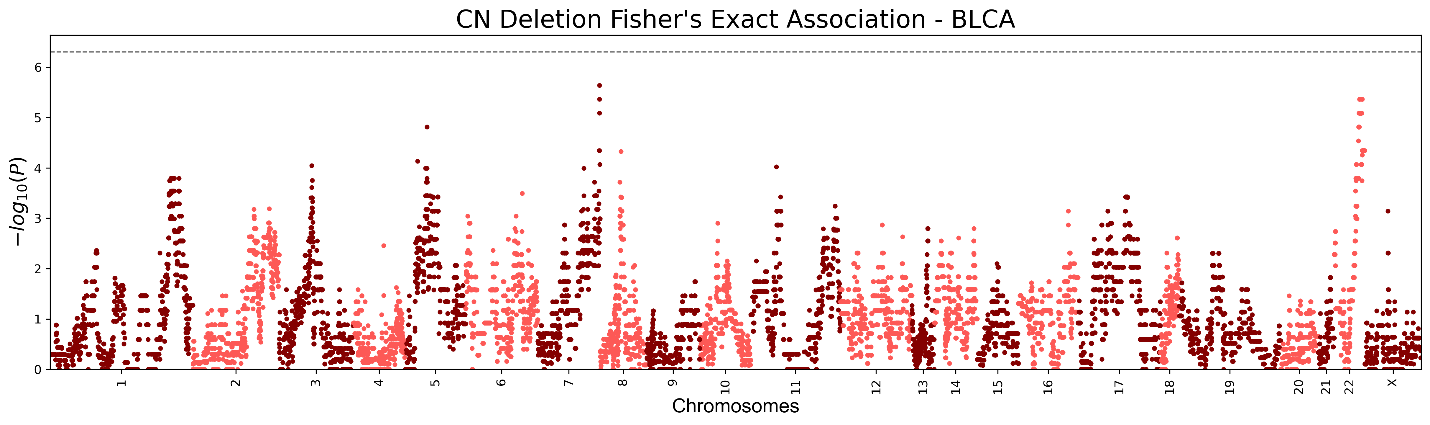

**C**

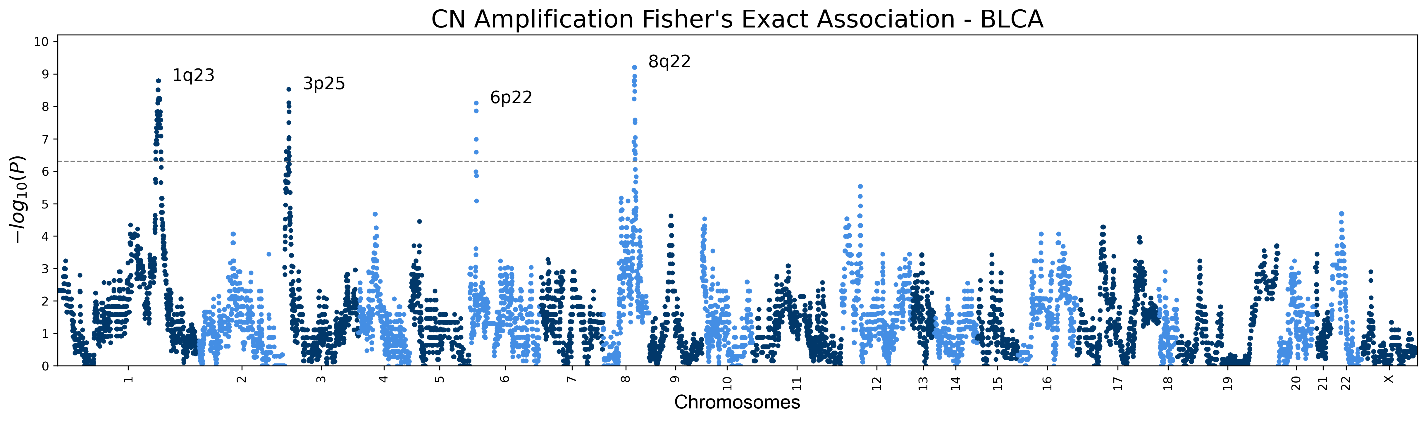

**Figure S3. BRCA Manhattan plots**

Genic Manhattan plot over Fisher’s exact test p-values between BRCA chromoanagenesis samples and non-chromoanagenesis samples. **(a)** Manhattan CNA (deletion or amplification) plot. **(b)** Manhattan Deletion plot. **(c)** Manhattan Amplification plot.

**a**
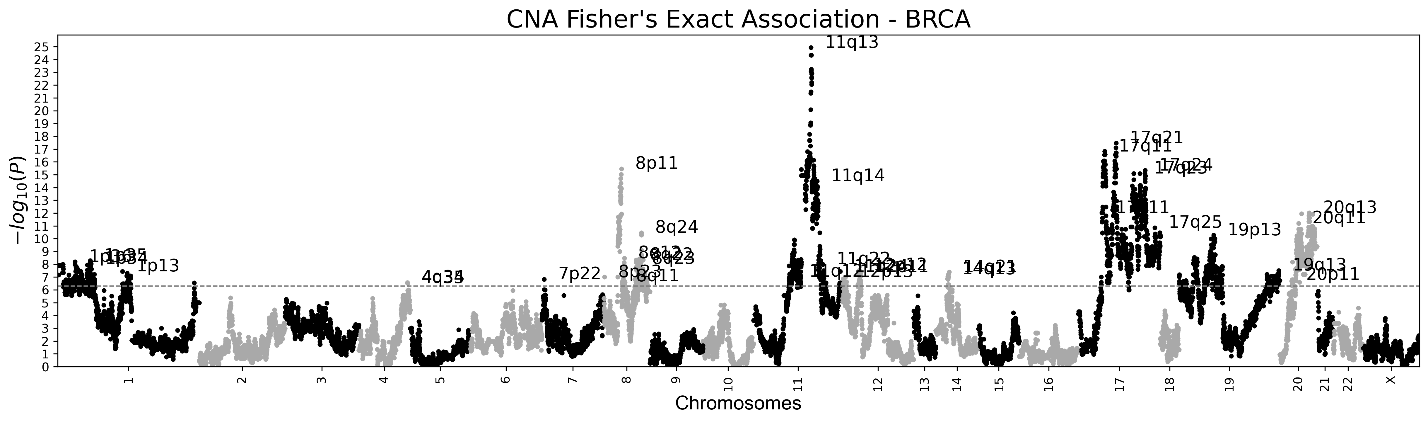

**b**
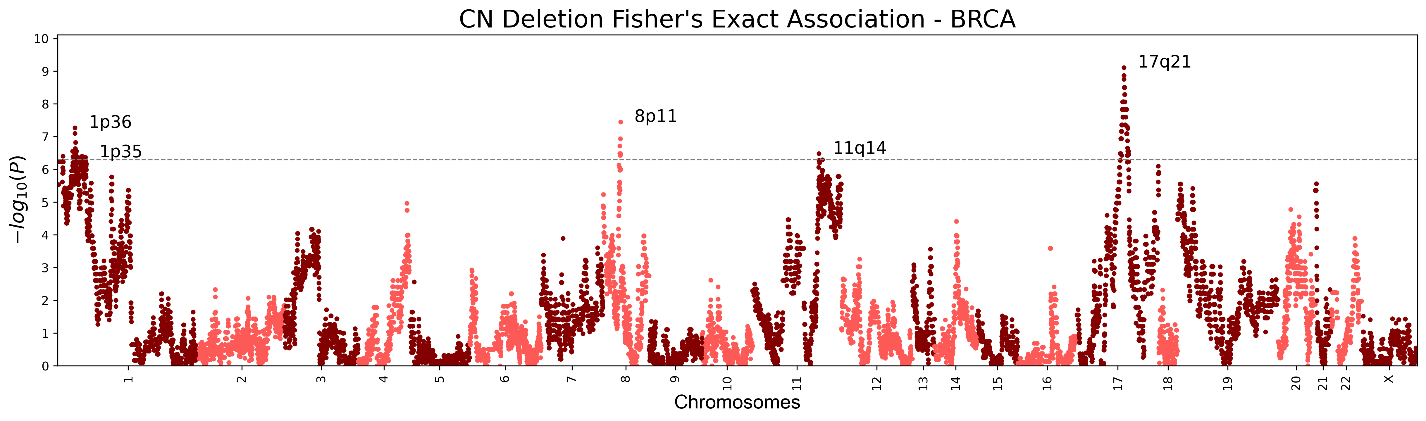

**c**
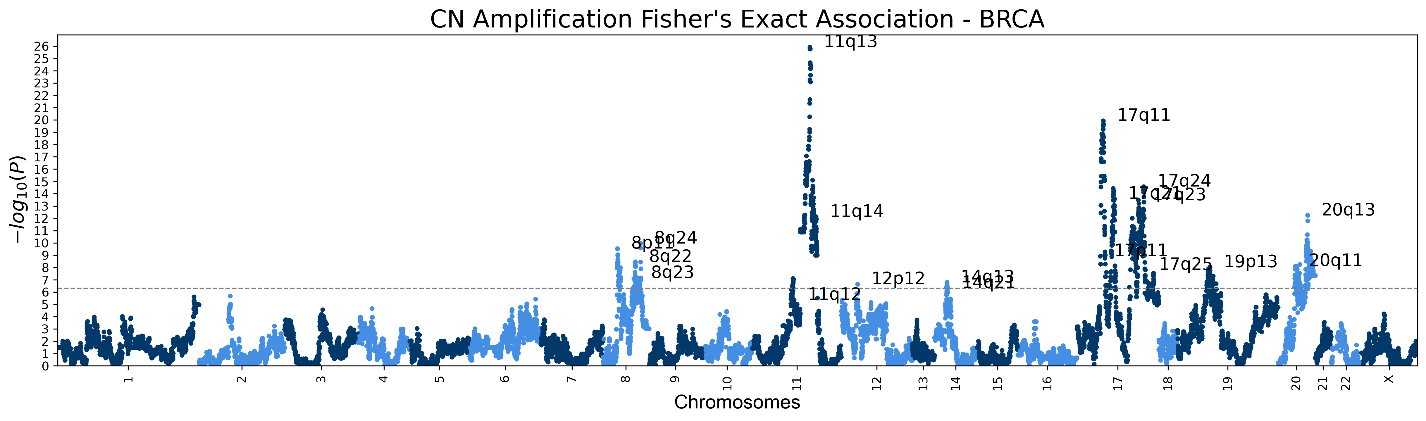

**Figure S4. CESC Manhattan plots**

Genic Manhattan plot over Fisher’s exact test p-values between CESC chromoanagenesis samples and non-chromoanagenesis samples. **(a)** Manhattan CNA (deletion or amplification) plot. **(b)** Manhattan Deletion plot. **(c)** Manhattan Amplification plot.

**a**
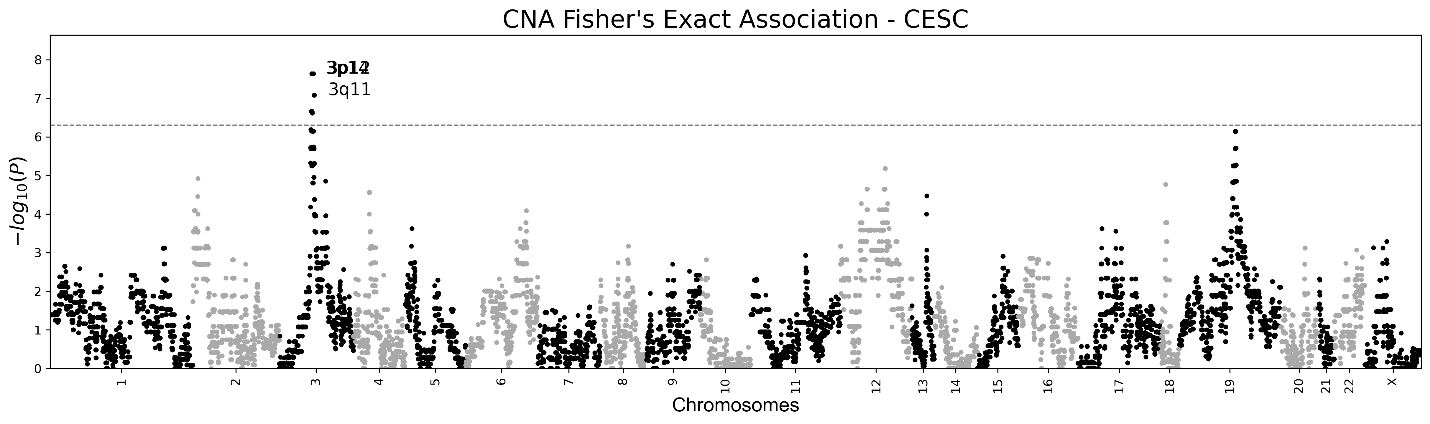

**b**
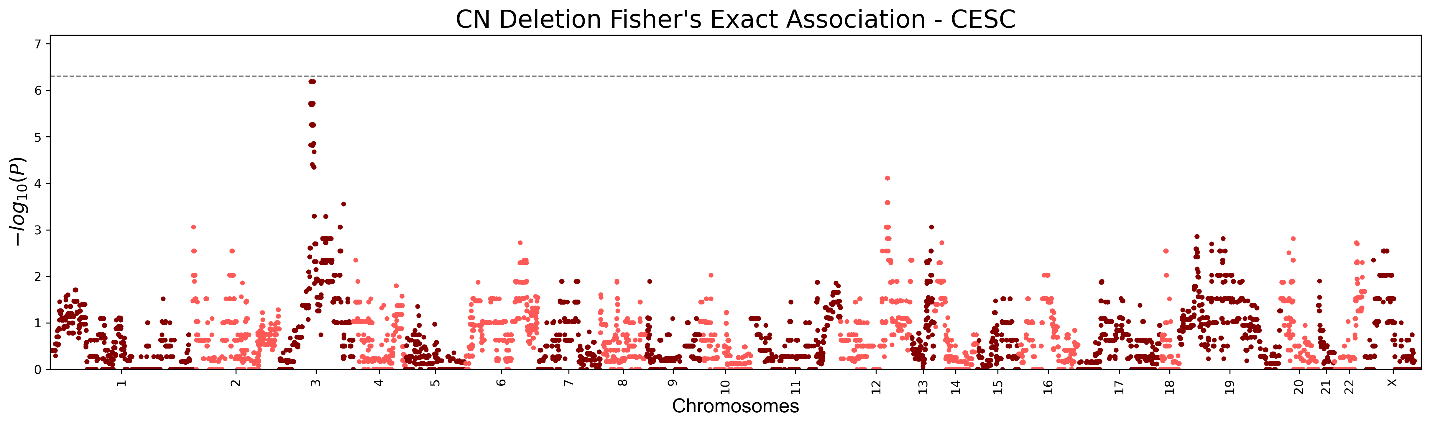

**c**
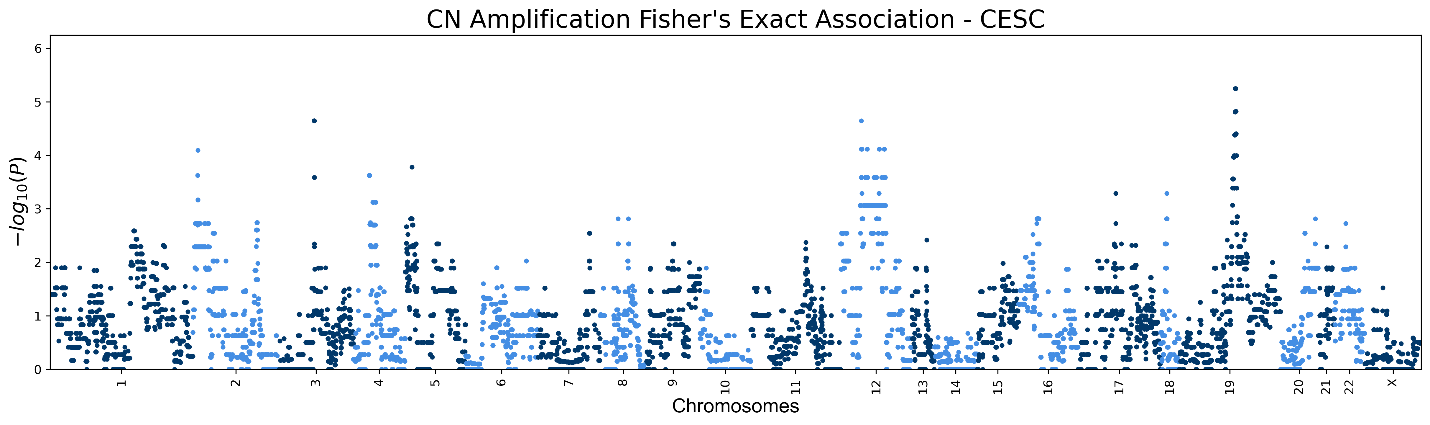

**Figure S5. COAD Manhattan plots**

Genic Manhattan plot over Fisher’s exact test p-values between COAD chromoanagenesis samples and non-chromoanagenesis samples. **(a)** Manhattan CNA (deletion or amplification) plot. **(b)** Manhattan Deletion plot. **(c)** Manhattan Amplification plot.

**a**
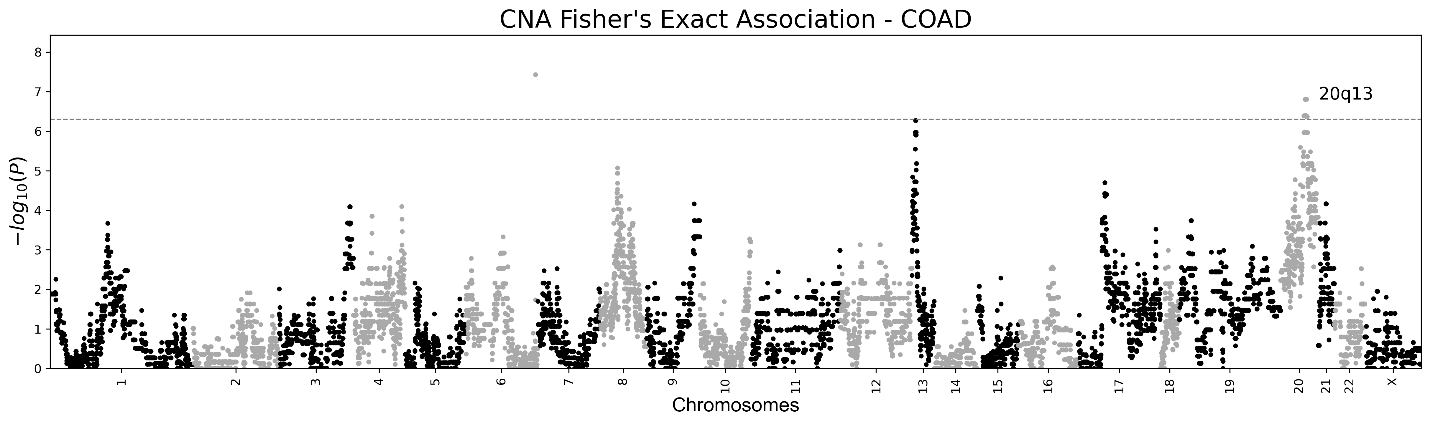

**b**
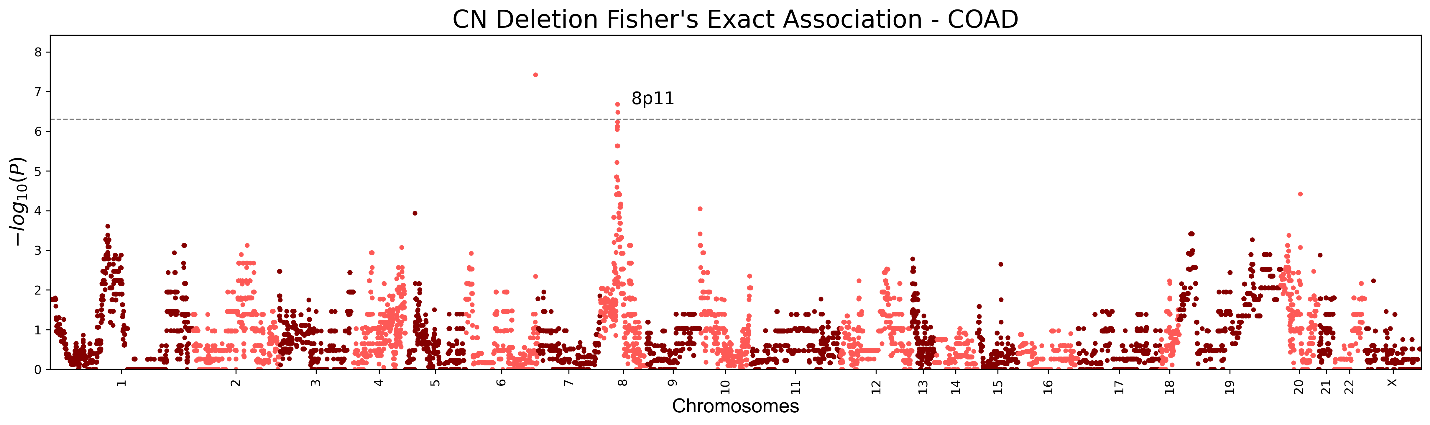

**c**
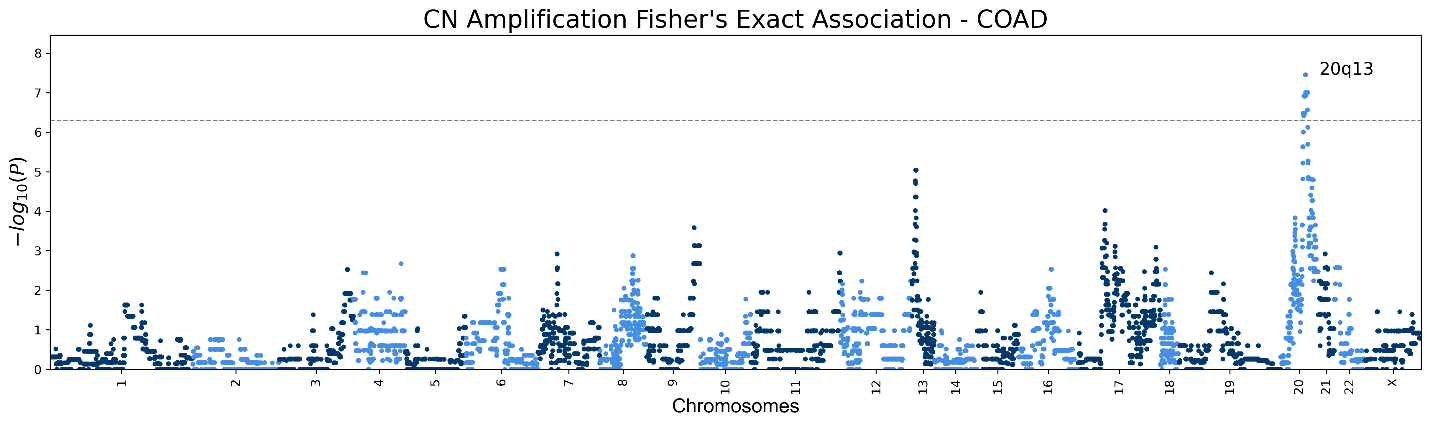

**Figure S6. ESCA Manhattan plots**

Genic Manhattan plot over Fisher’s exact test p-values between ESCA chromoanagenesis samples and non-chromoanagenesis samples. **(a)** Manhattan CNA (deletion or amplification) plot. **(b)** Manhattan Deletion plot. **(c)** Manhattan Amplification plot.

**a**
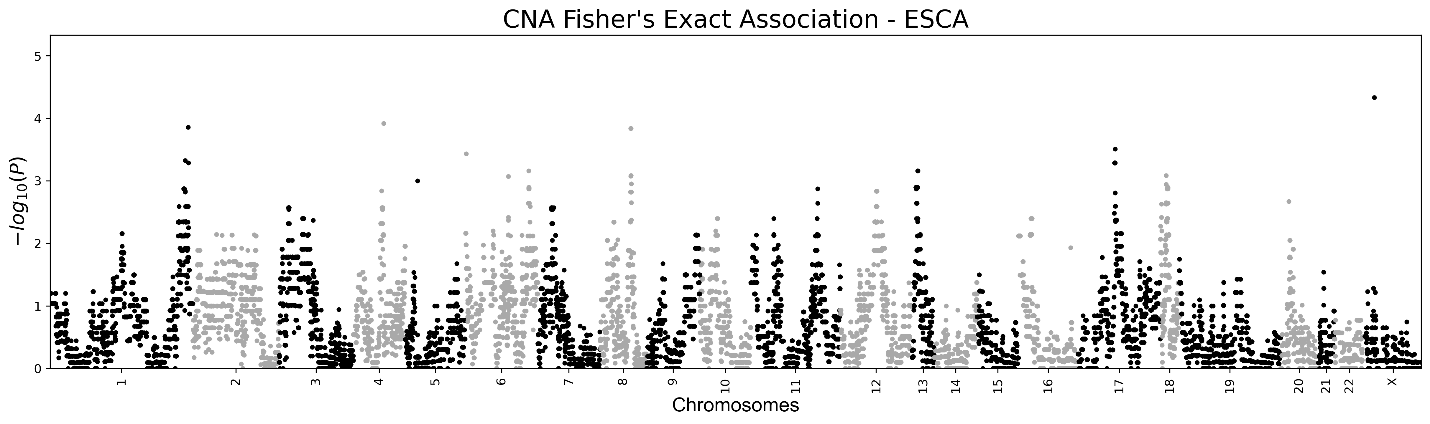

**b**
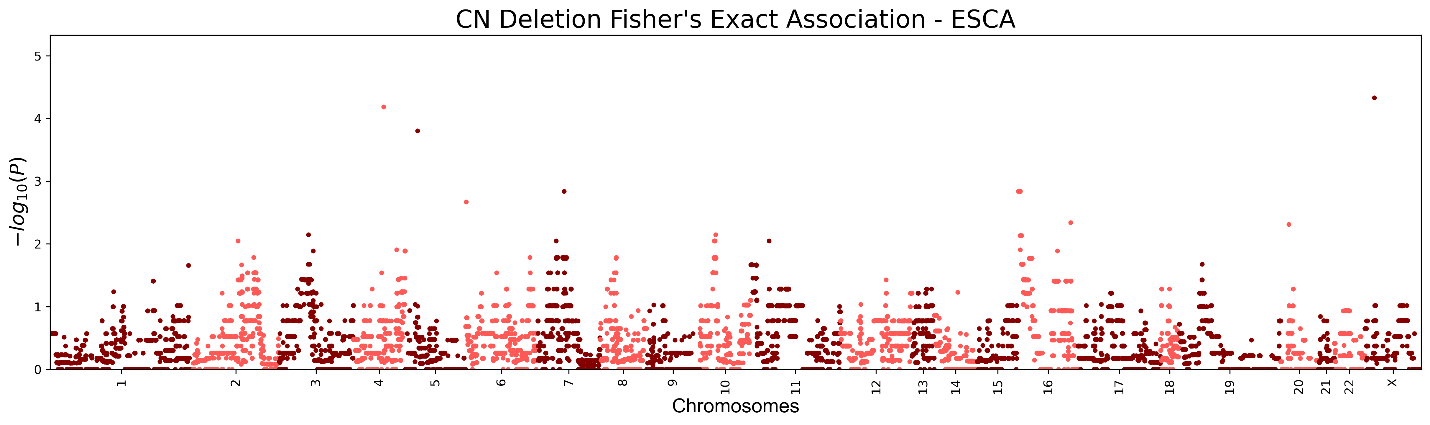

**c**
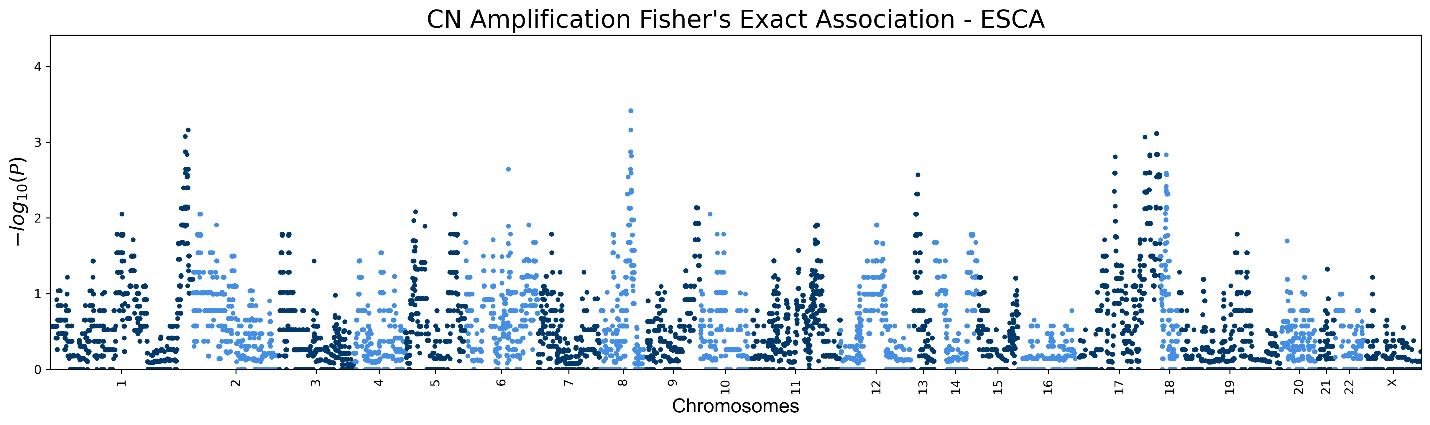

**Figure S7. GBM Manhattan plots**

Genic Manhattan plot over Fisher’s exact test p-values between GBM chromoanagenesis samples and non-chromoanagenesis samples. **(a)** Manhattan CNA (deletion or amplification) plot. **(b)** Manhattan Deletion plot. **(c)** Manhattan Amplification plot.

**a**
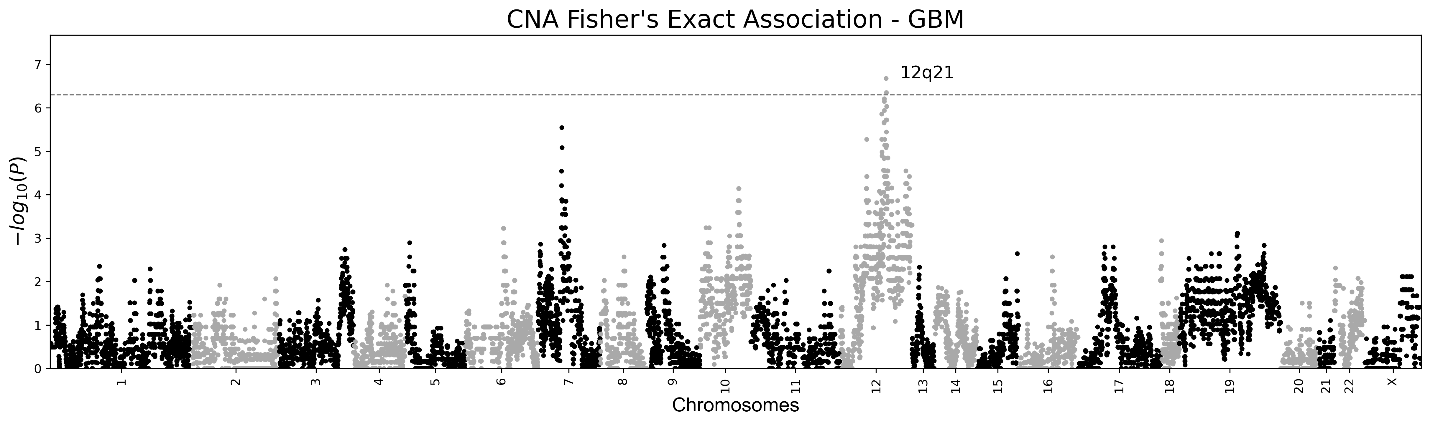

**b**
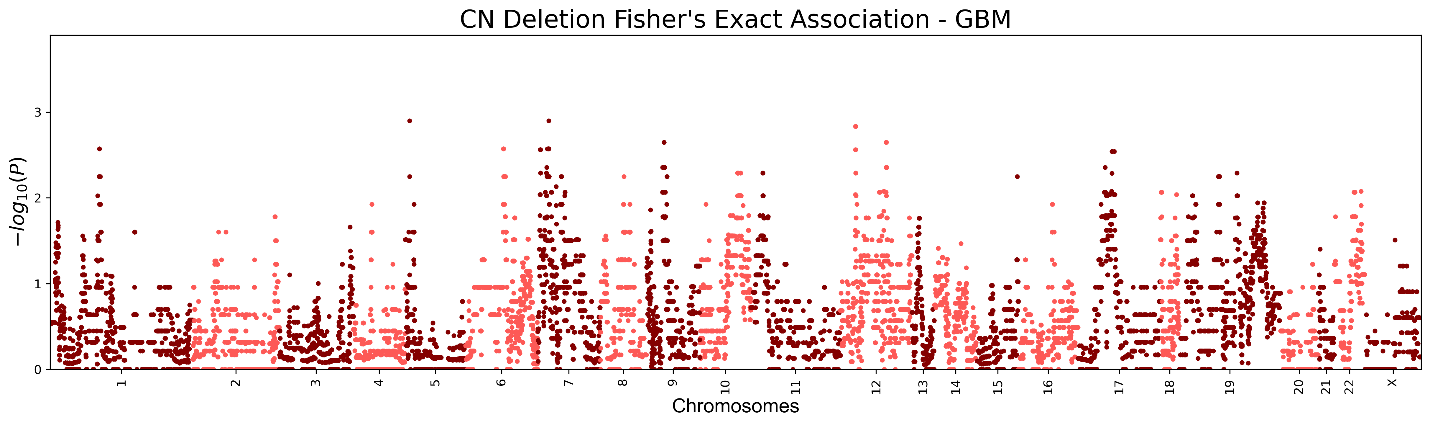

**c**
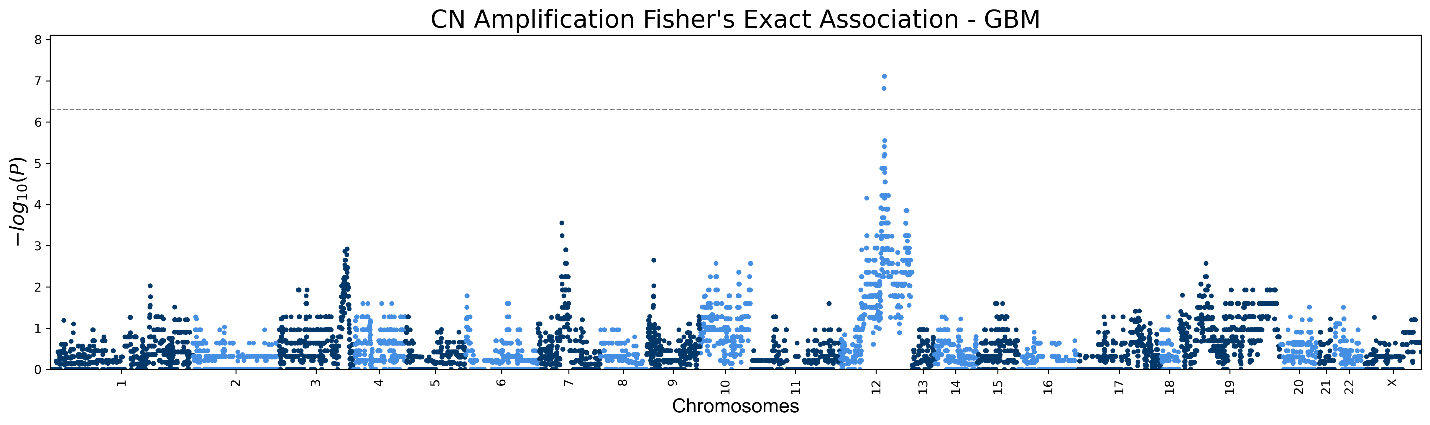

**Figure S8. HNSC Manhattan plots**

Genic Manhattan plot over Fisher’s exact test p-values between HNSC chromoanagenesis samples and non-chromoanagenesis samples. **(a)** Manhattan CNA (deletion or amplification) plot. **(b)** Manhattan Deletion plot. **(c)** Manhattan Amplification plot.

**a**
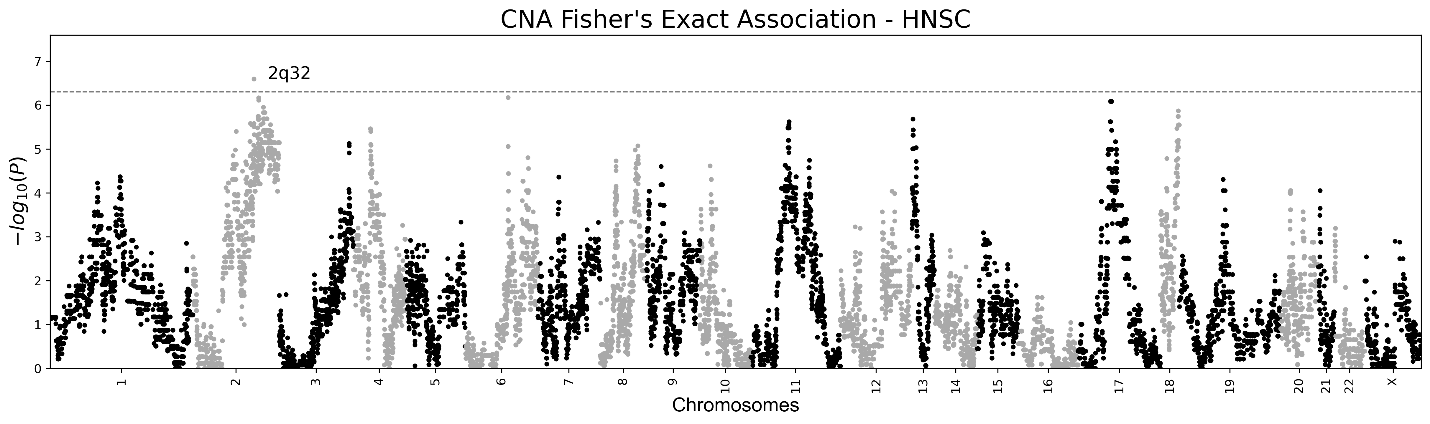

**b**
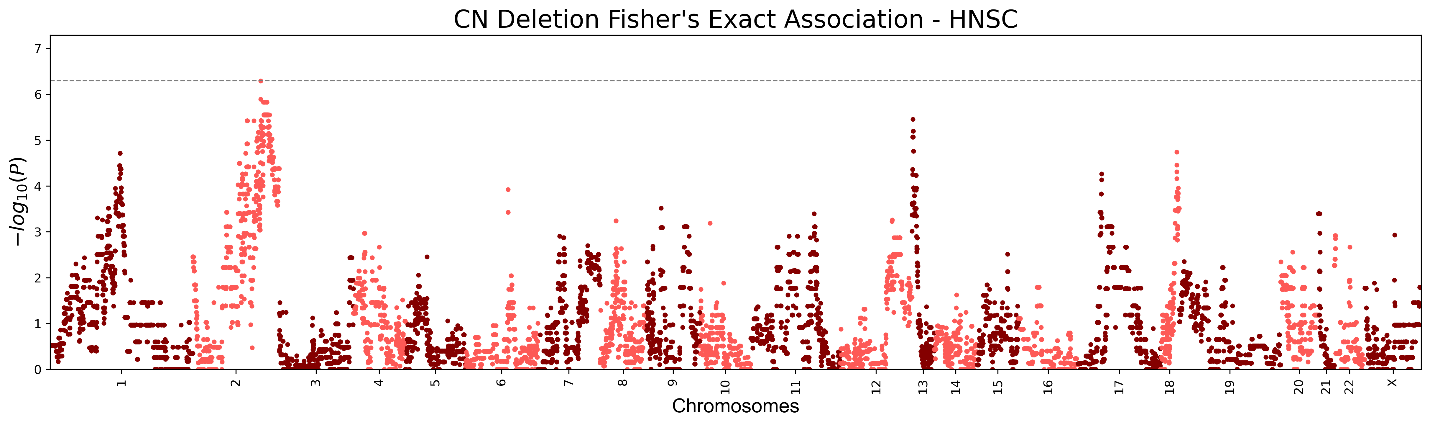

**c**
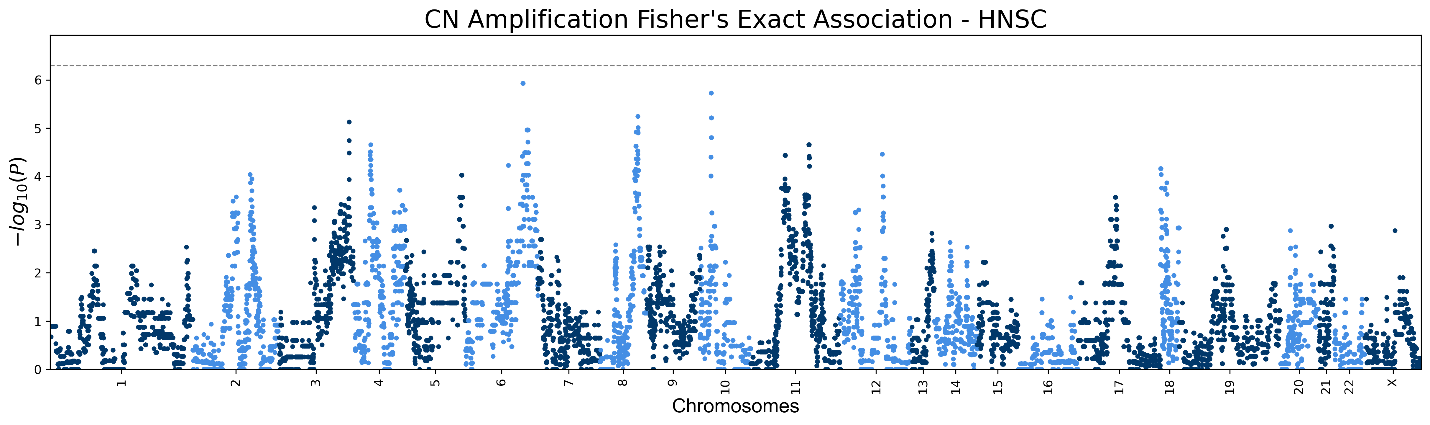

**Figure S9. KIRC Manhattan plots**

Genic Manhattan plot over Fisher’s exact test p-values between KIRC chromoanagenesis samples and non-chromoanagenesis samples. **(a)** Manhattan CNA (deletion or amplification) plot. **(b)** Manhattan Deletion plot. **(c)** Manhattan Amplification plot.

**a**
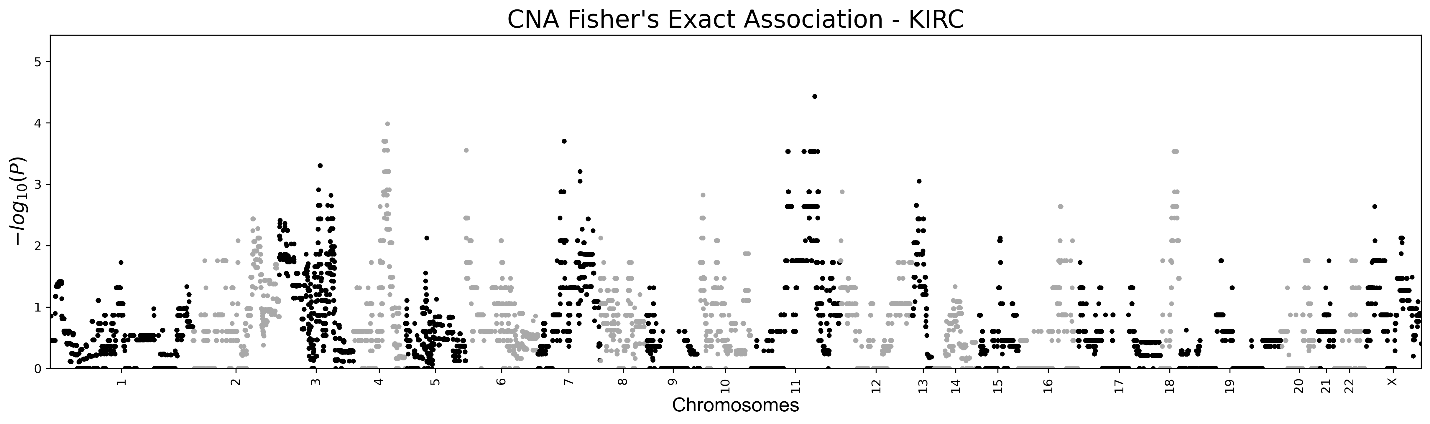

**b**
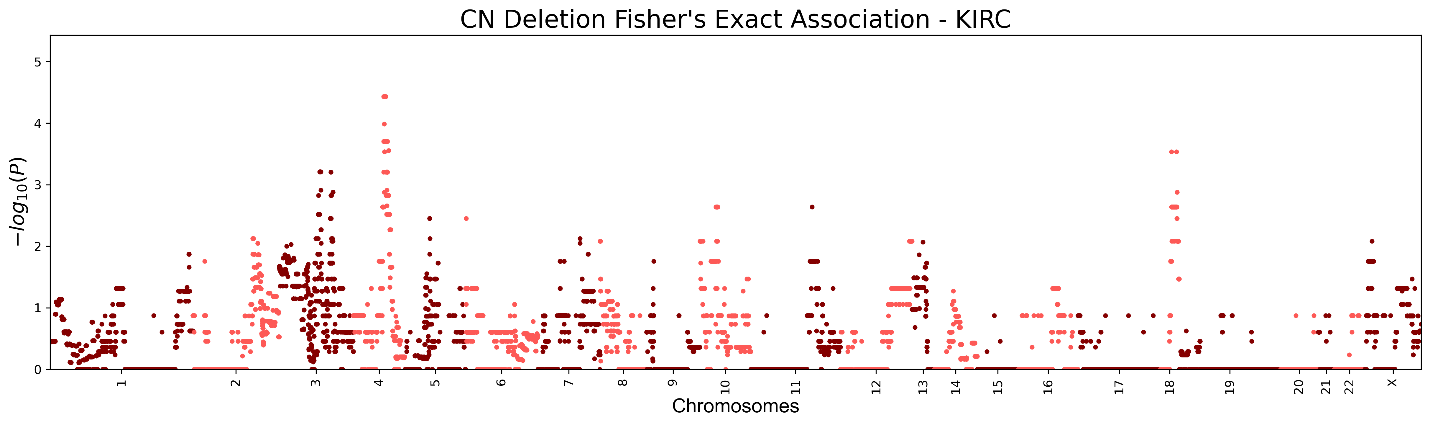

**c**
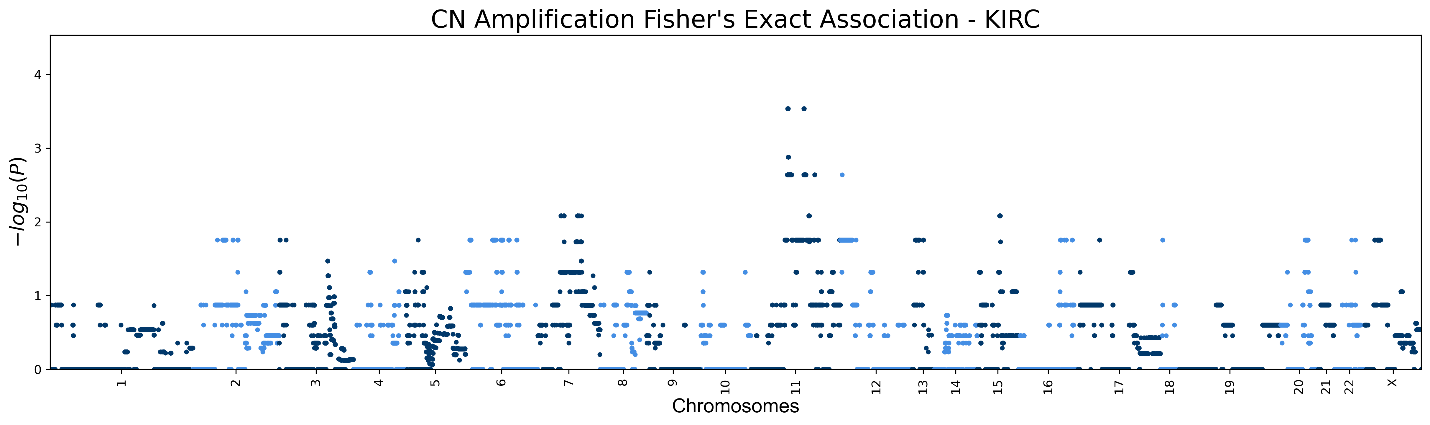

**Figure S10. LIHC Manhattan plots**

Genic Manhattan plot over Fisher’s exact test p-values between LIHC chromoanagenesis samples and non-chromoanagenesis samples. **(a)** Manhattan CNA (deletion or amplification) plot. **(b)** Manhattan Deletion plot. **(c)** Manhattan Amplification plot.

**a**
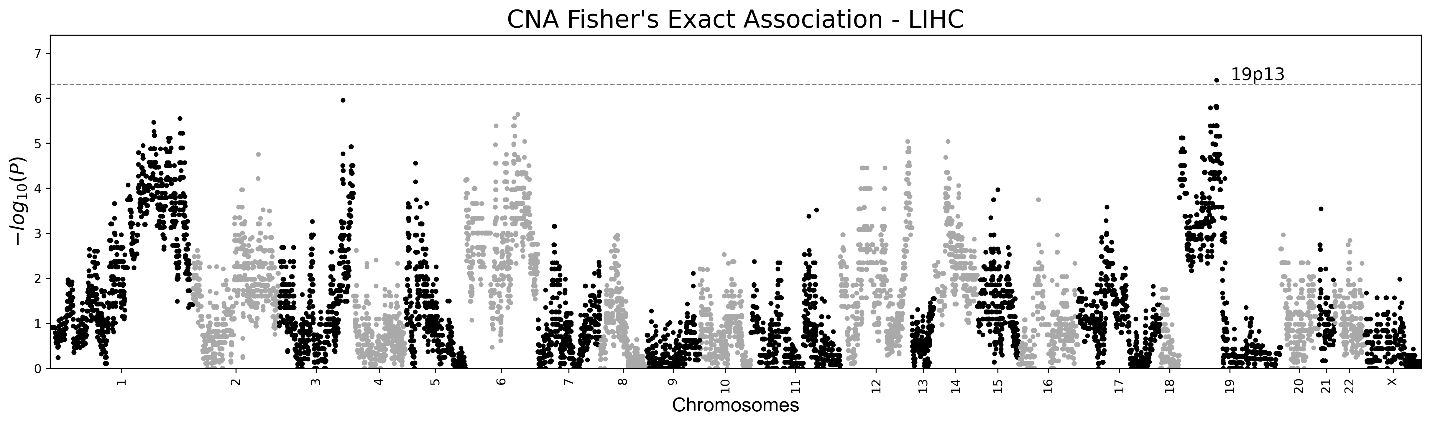

**b**
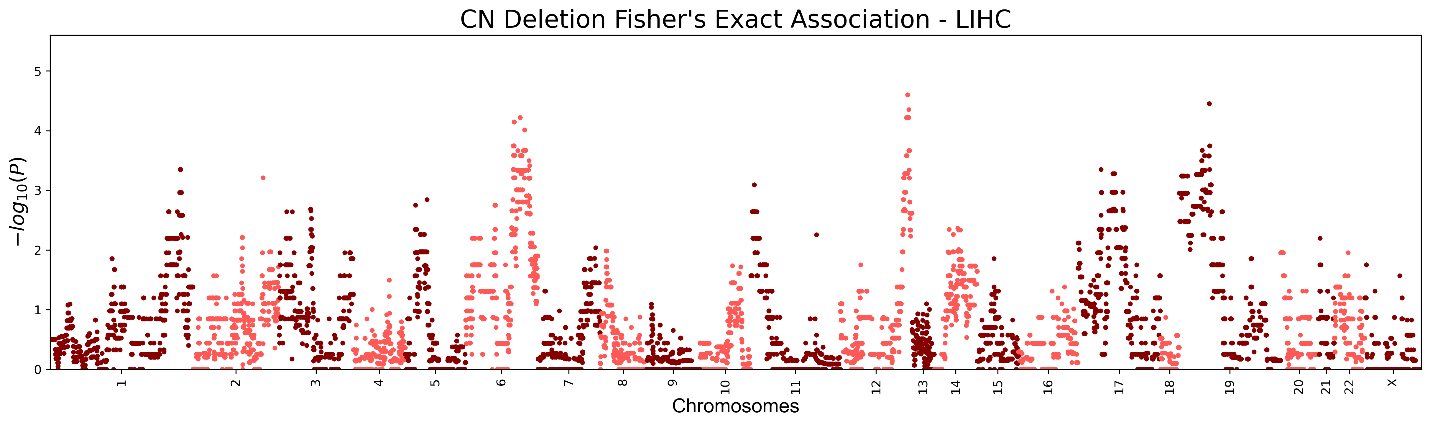

**c**
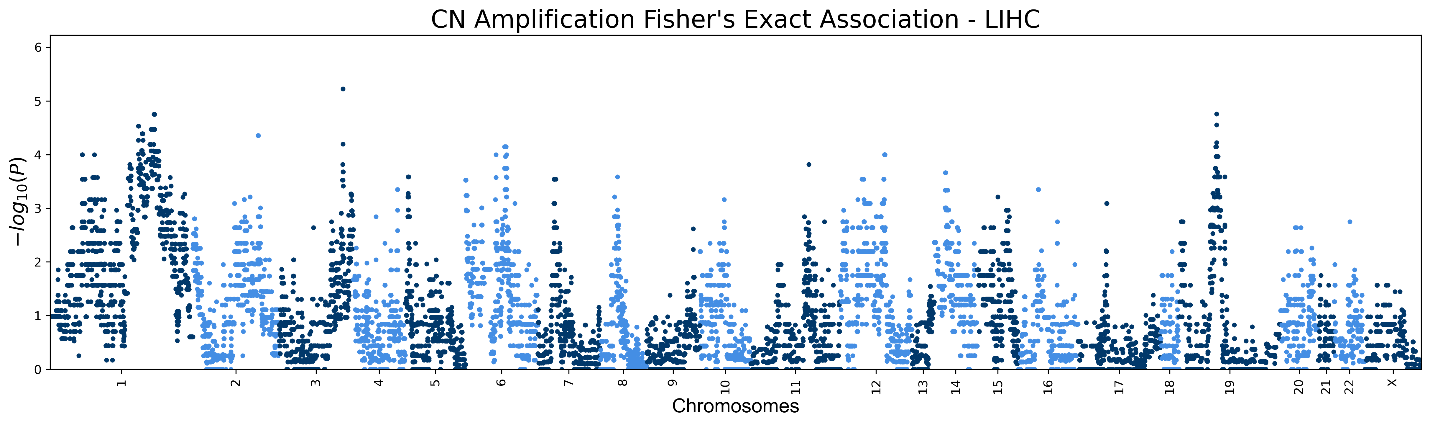

**Figure S11. LUAD Manhattan plots**

Genic Manhattan plot over Fisher’s exact test p-values between LUAD chromoanagenesis samples and non-chromoanagenesis samples. **(a)** Manhattan CNA (deletion or amplification) plot. **(b)** Manhattan Deletion plot. **(c)** Manhattan Amplification plot.

**a**
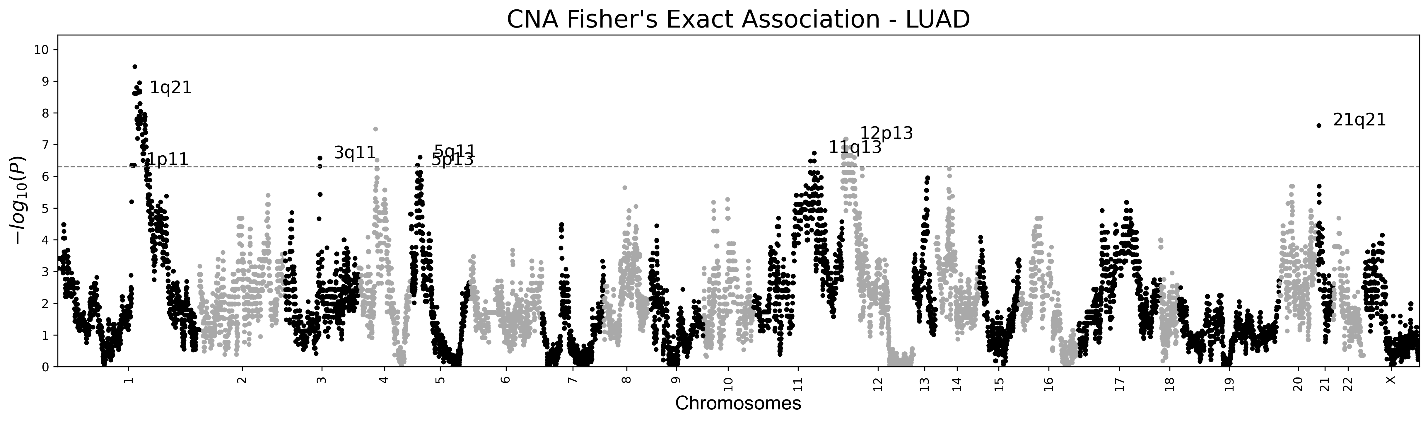

**b**
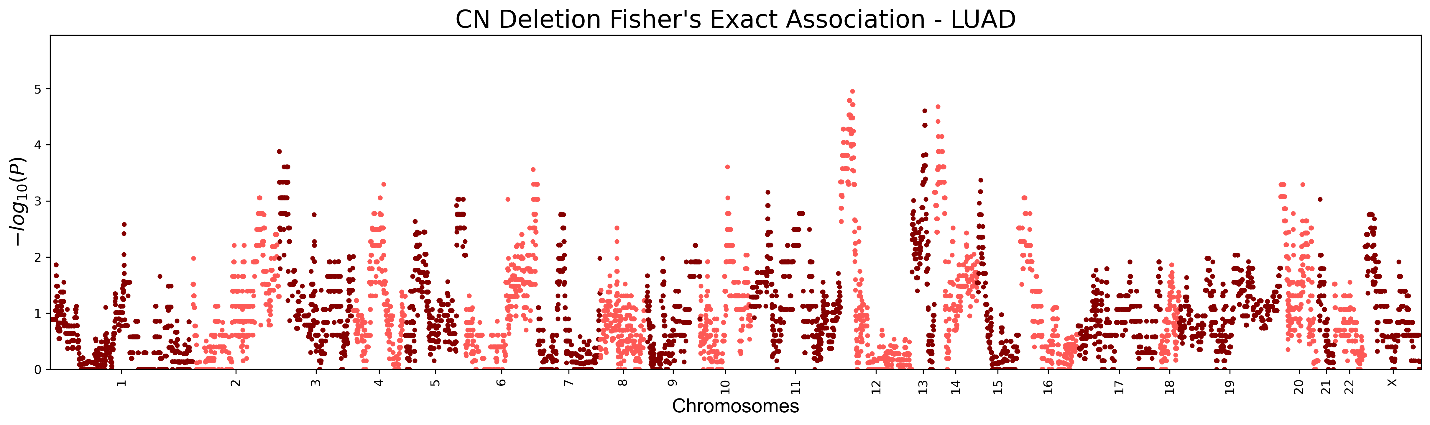

**c**
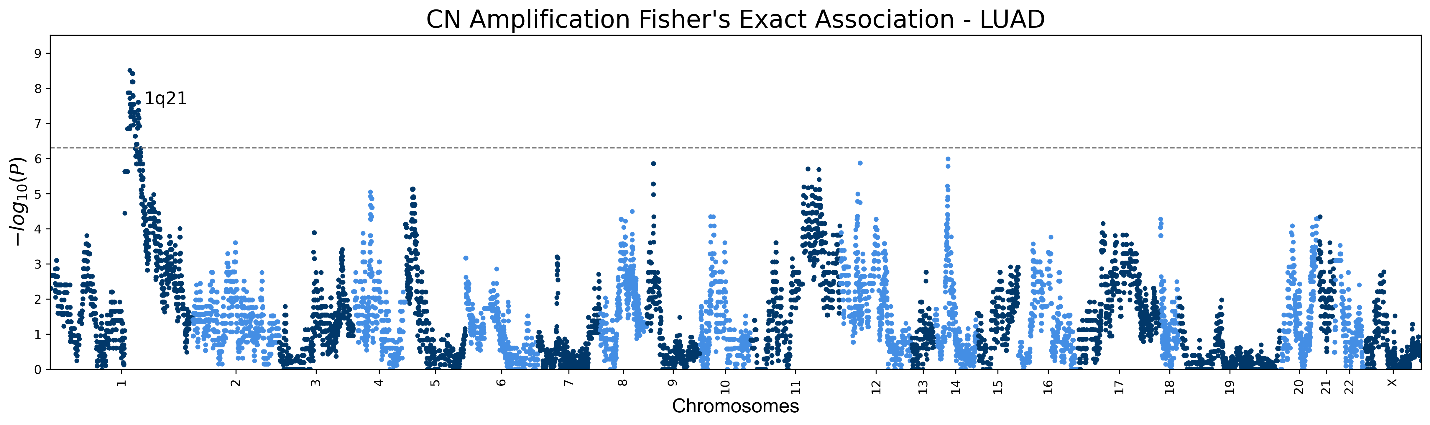

**Figure S12. LUSC Manhattan plots**

Genic Manhattan plot over Fisher’s exact test p-values between LUSC chromoanagenesis samples and non-chromoanagenesis samples. **(a)** Manhattan CNA (deletion or amplification) plot. **(b)** Manhattan Deletion plot. **(c)** Manhattan Amplification plot.

**a**
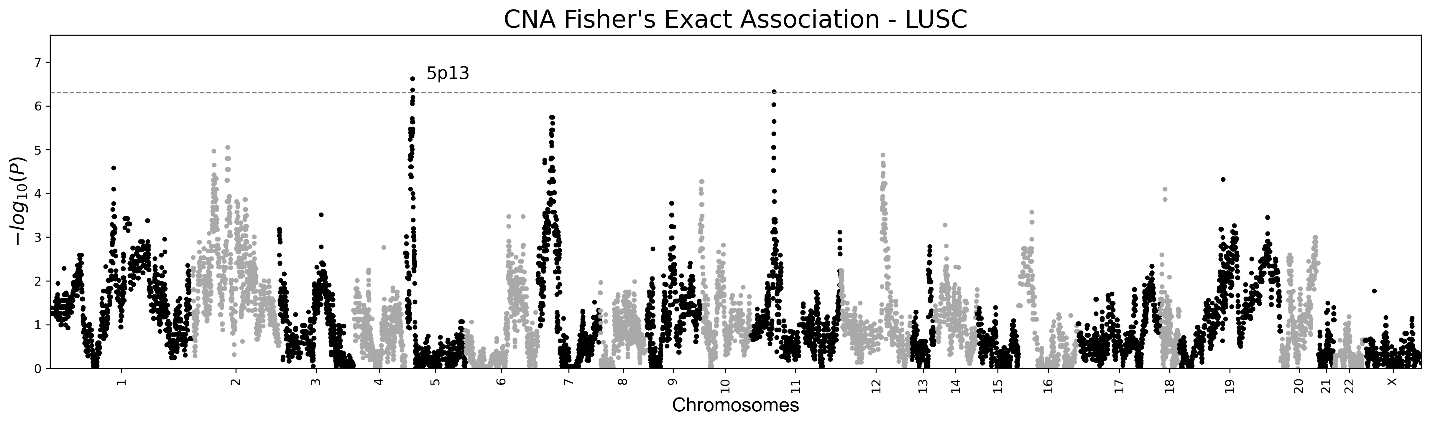

**b**
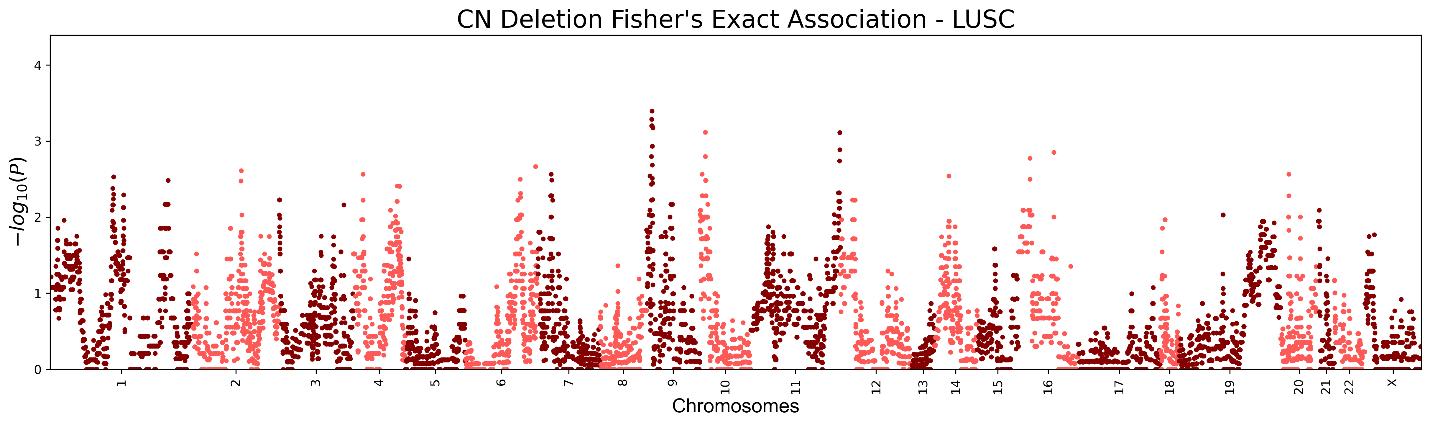

**c**
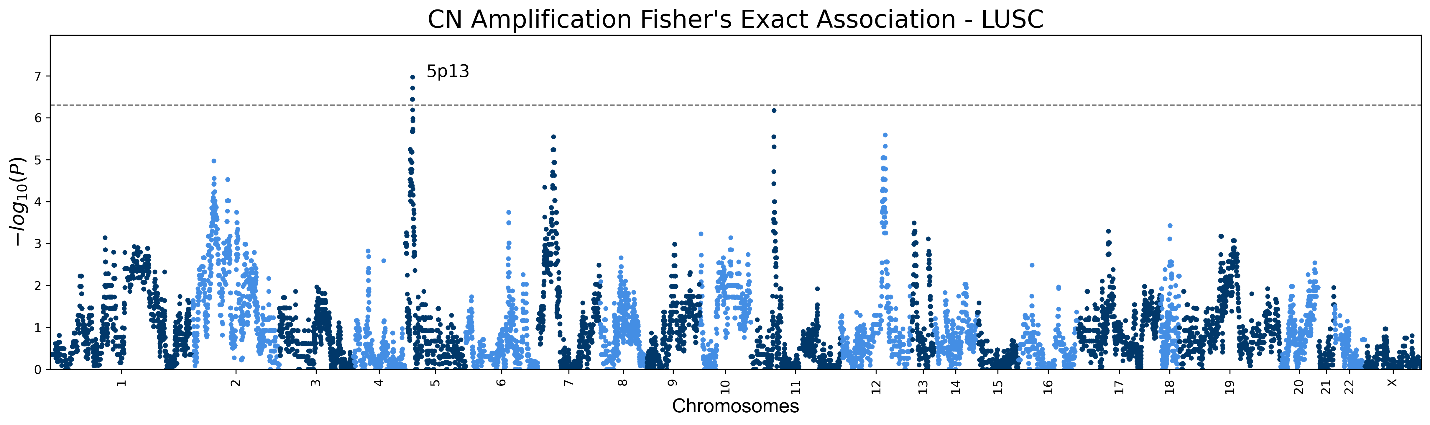

**Figure S13. OV Manhattan plots**

Genic Manhattan plot over Fisher’s exact test p-values between OV chromoanagenesis samples and non-chromoanagenesis samples. **(a)** Manhattan CNA (deletion or amplification) plot. **(b)** Manhattan Deletion plot. **(c)** Manhattan Amplification plot.

**a**
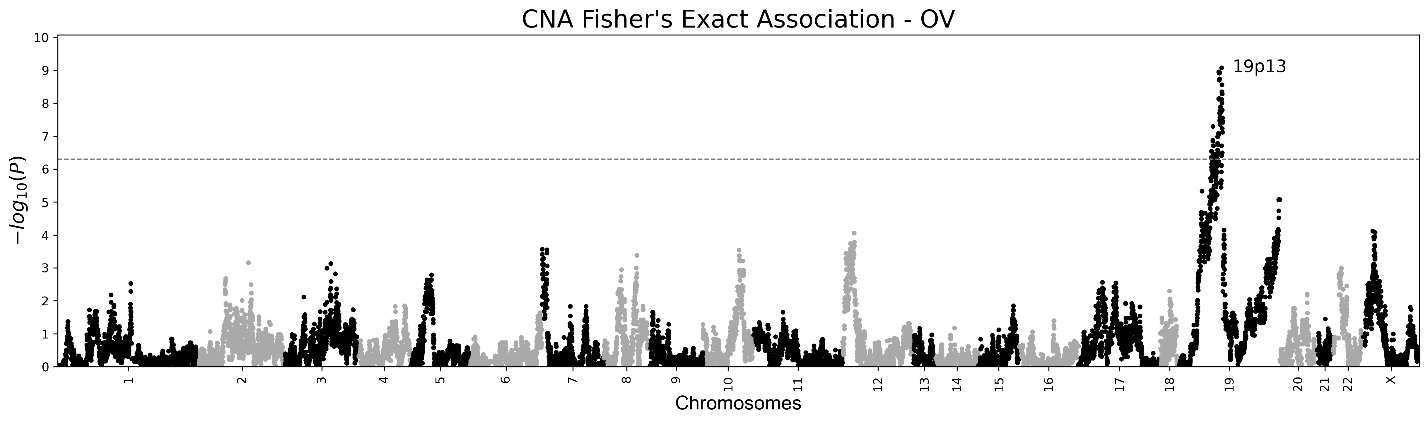

**b**

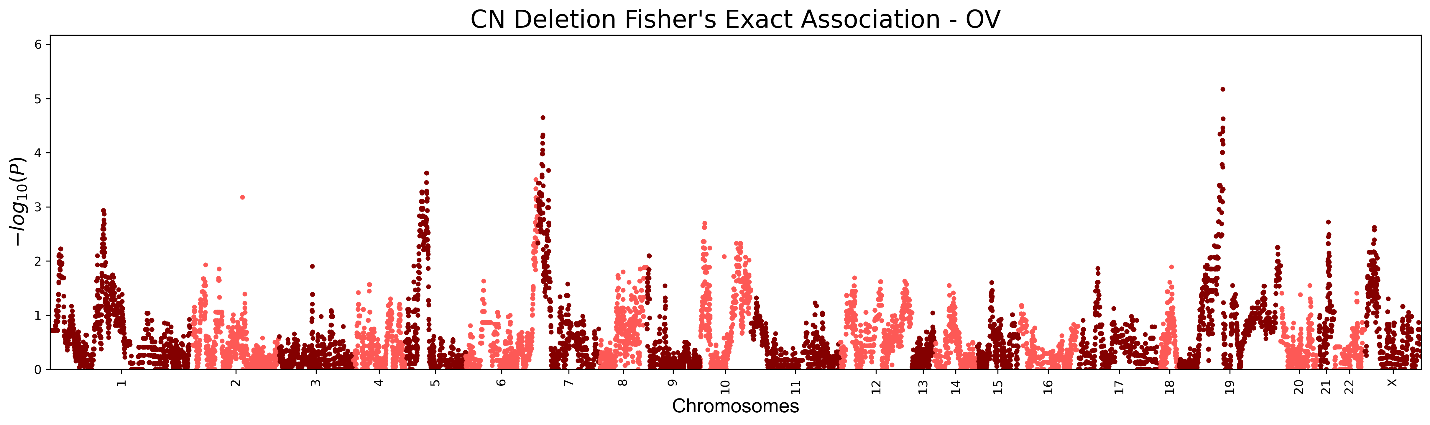

**c**
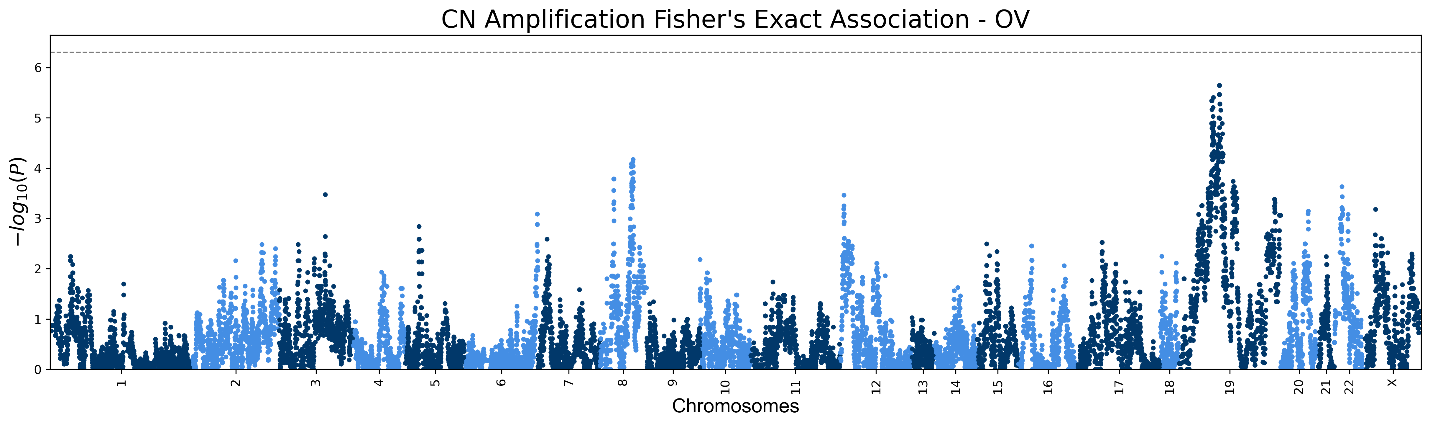

**Figure S14. PAAD Manhattan plots**

Genic Manhattan plot over Fisher’s exact test p-values between PAAD chromoanagenesis samples and non-chromoanagenesis samples. **(a)** Manhattan CNA (deletion or amplification) plot. **(b)** Manhattan Deletion plot. **(c)** Manhattan Amplification plot.

**a**
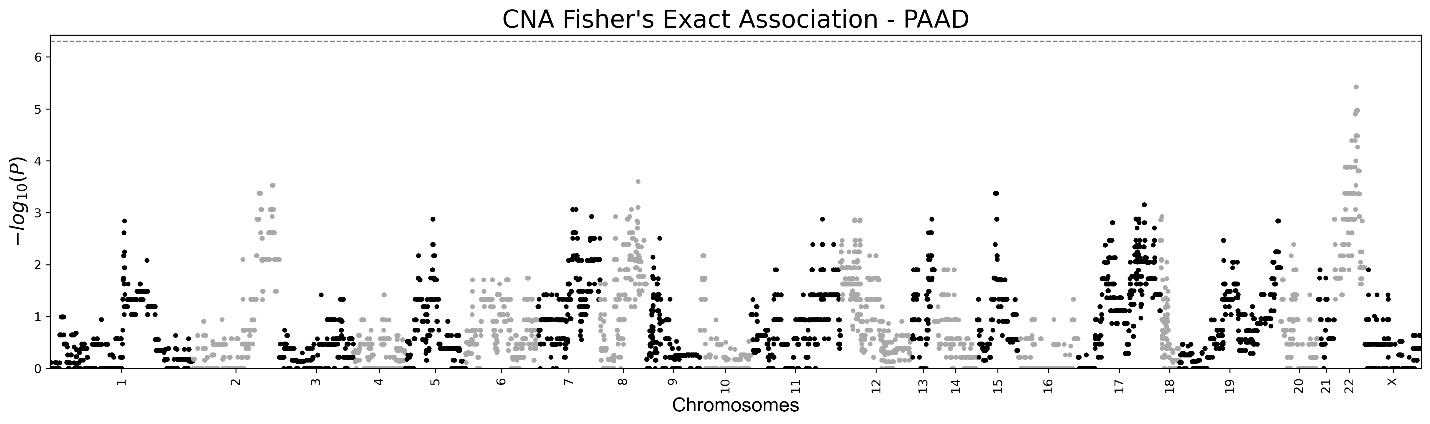

**b**
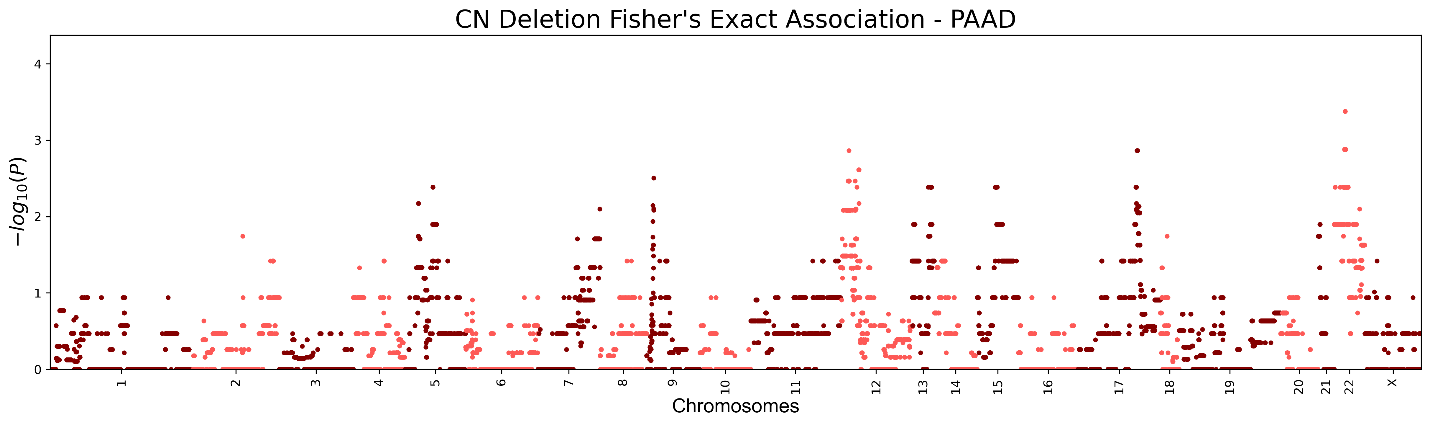

**c**
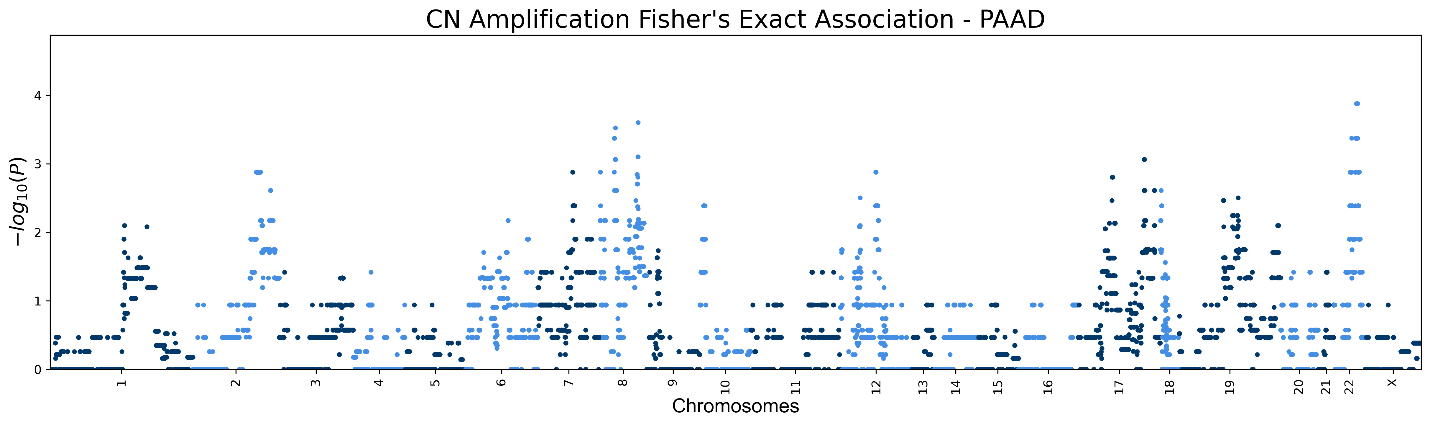

**Figure S15. PRAD Manhattan plots**

Genic Manhattan plot over Fisher’s exact test p-values between PRAD chromoanagenesis samples and non-chromoanagenesis samples. **(a)** Manhattan CNA (deletion or amplification) plot. **(b)** Manhattan Deletion plot. **(c)** Manhattan Amplification plot.

**a**
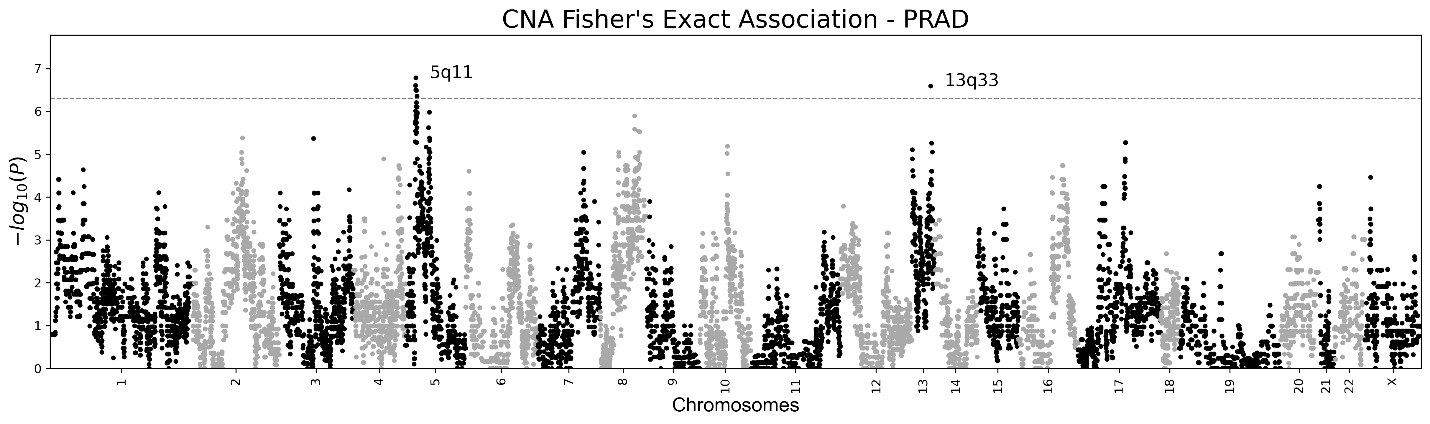

**b**
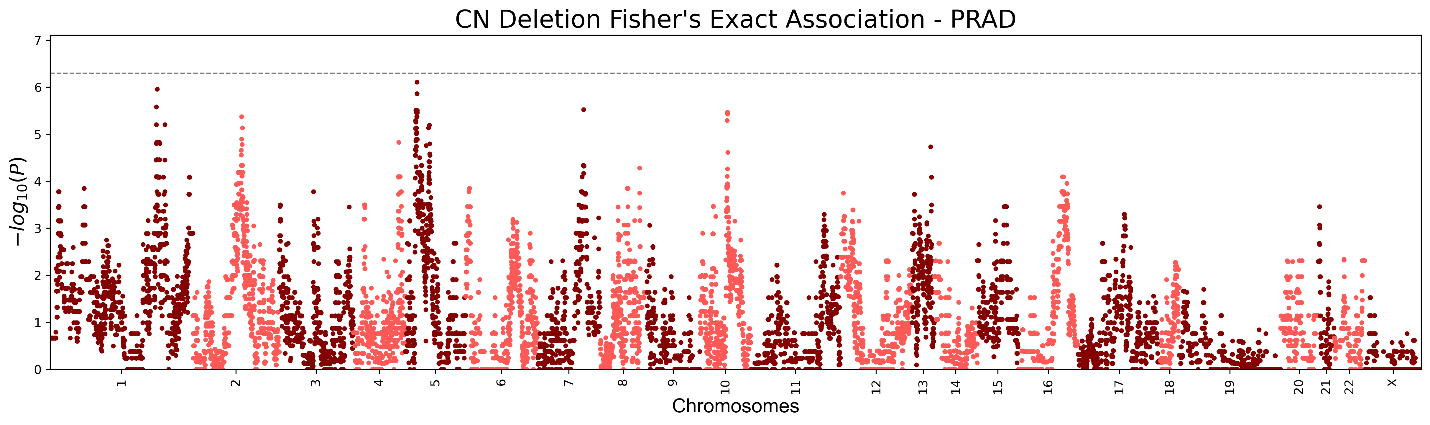

**c**
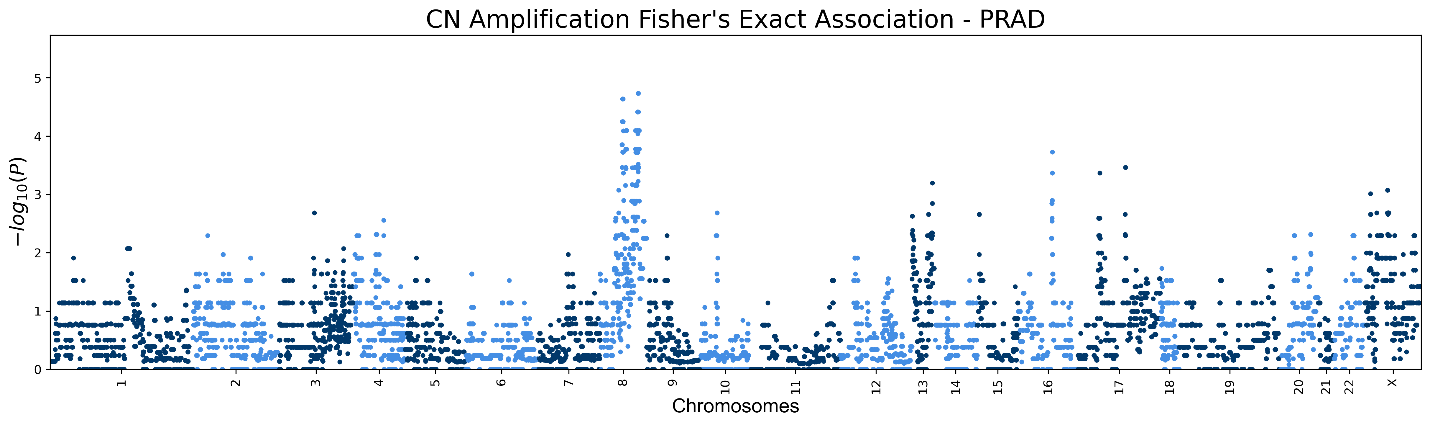

**Figure S16. READ Manhattan plots**

Genic Manhattan plot over Fisher’s exact test p-values between READ chromoanagenesis samples and non-chromoanagenesis samples. **(a)** Manhattan CNA (deletion or amplification) plot. **(b)** Manhattan Deletion plot. **(c)** Manhattan Amplification plot.

**a**
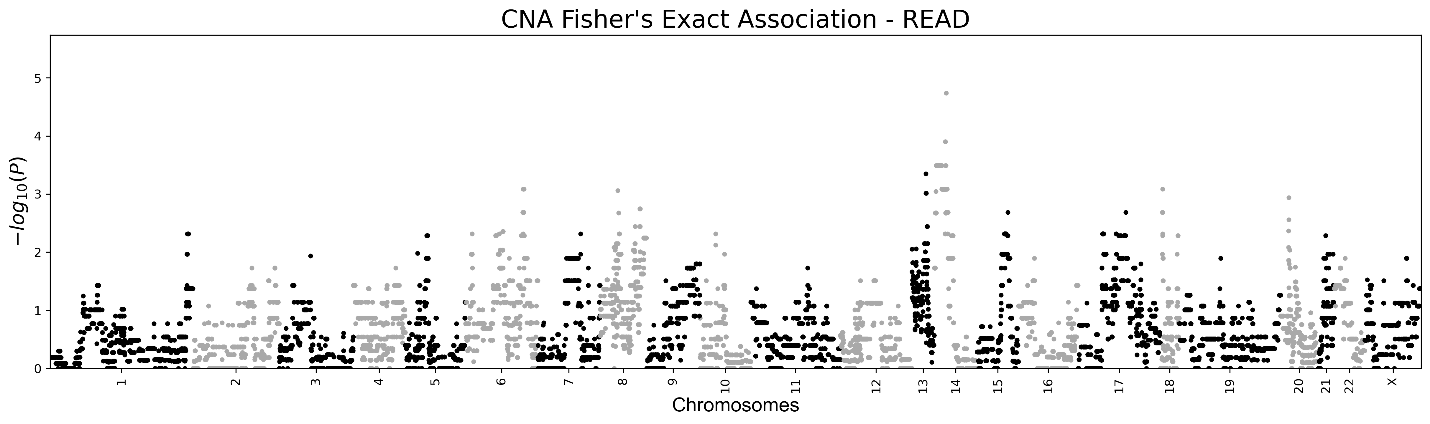

**b**
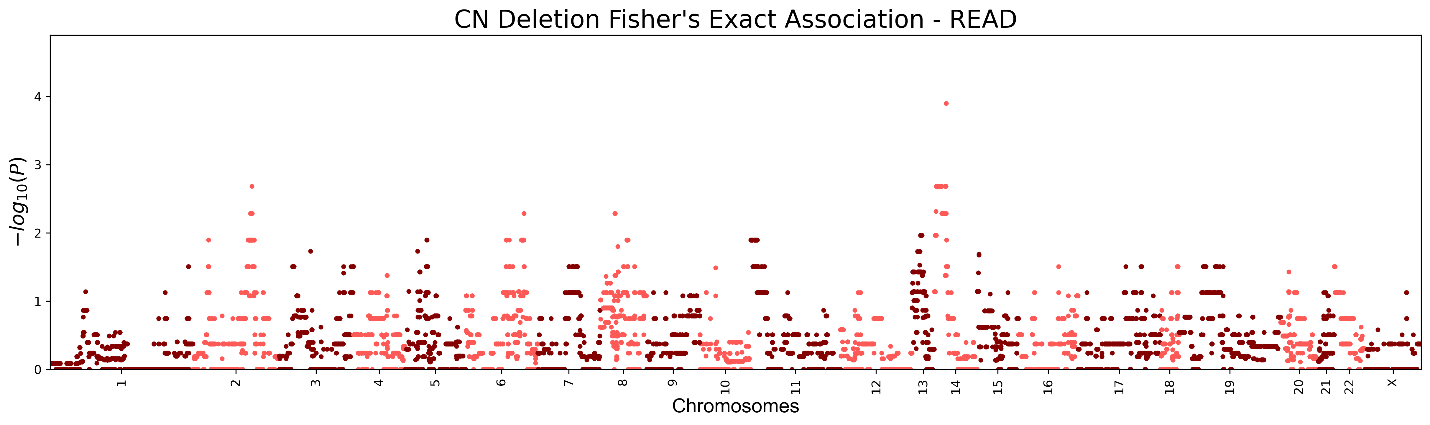

**c**
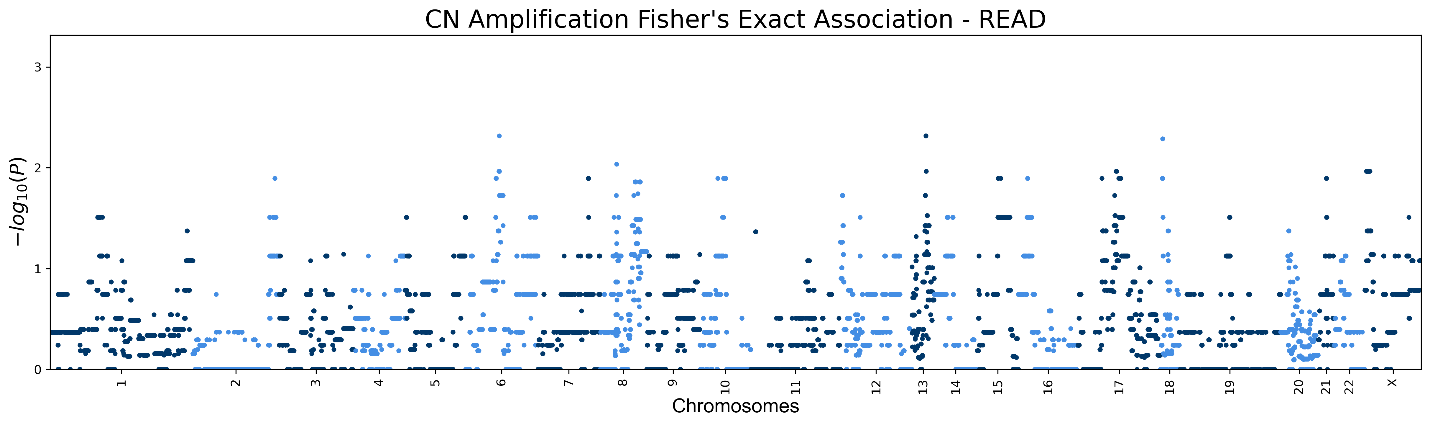

**Figure S17. SARC Manhattan plots**

Genic Manhattan plot over Fisher’s exact test p-values between SARC chromoanagenesis samples and non-chromoanagenesis samples. **(a)** Manhattan CNA (deletion or amplification) plot. **(b)** Manhattan Deletion plot. **(c)** Manhattan Amplification plot.

**a**
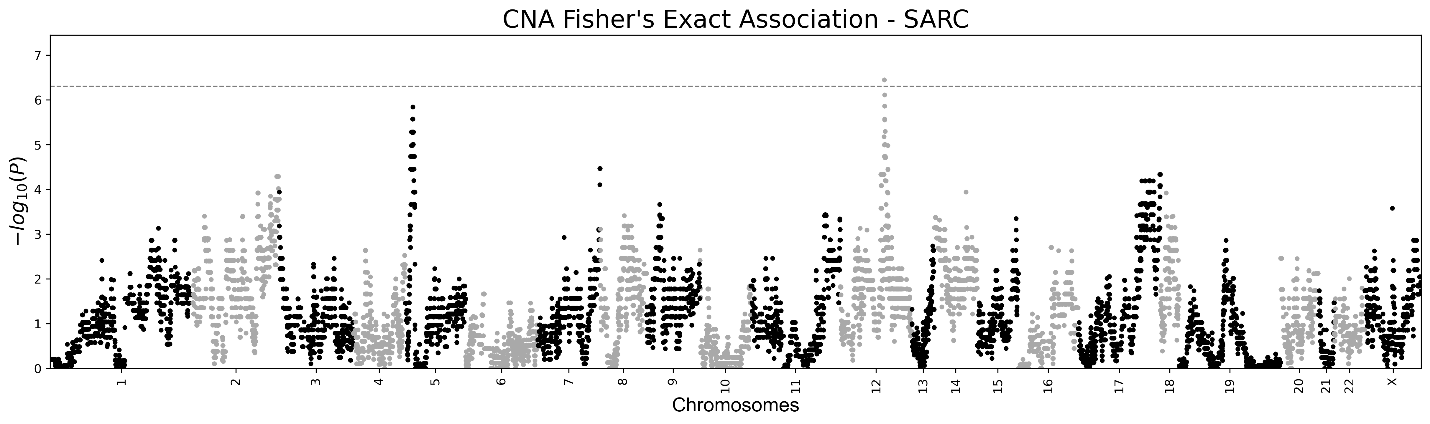

**b**
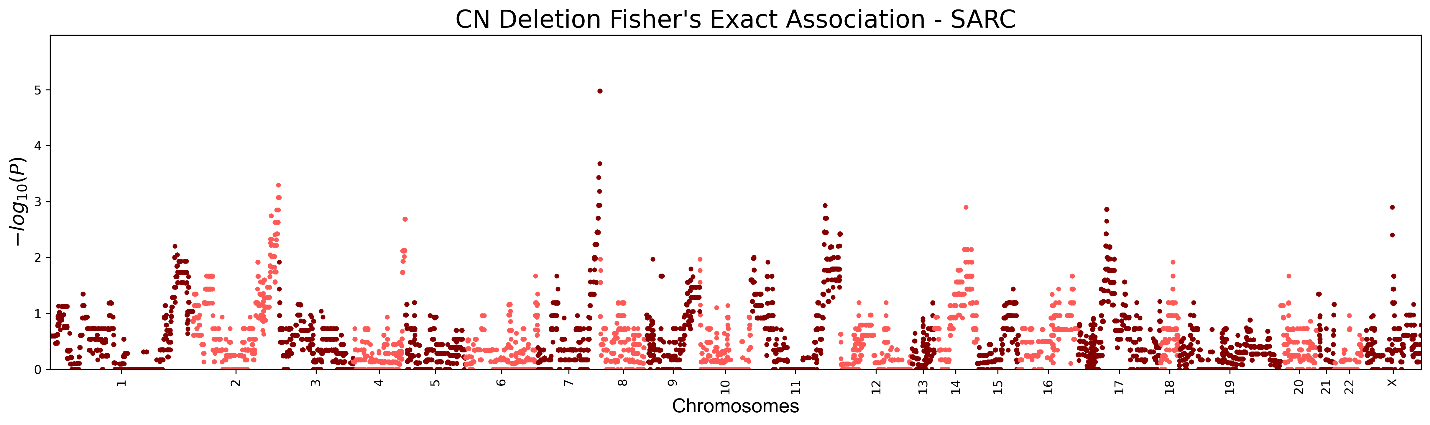

**c**
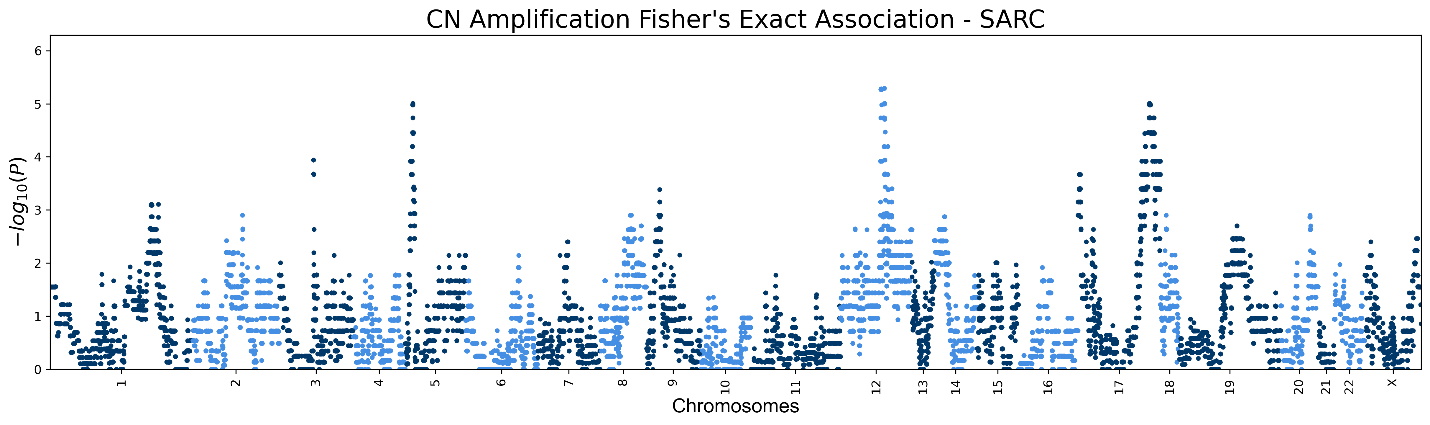

**Figure S18. SKCM Manhattan plots**

Genic Manhattan plot over Fisher’s exact test p-values between SKCM chromoanagenesis samples and non-chromoanagenesis samples. **(a)** Manhattan CNA (deletion or amplification) plot. **(b)** Manhattan Deletion plot. **(c)** Manhattan Amplification plot.

**a**
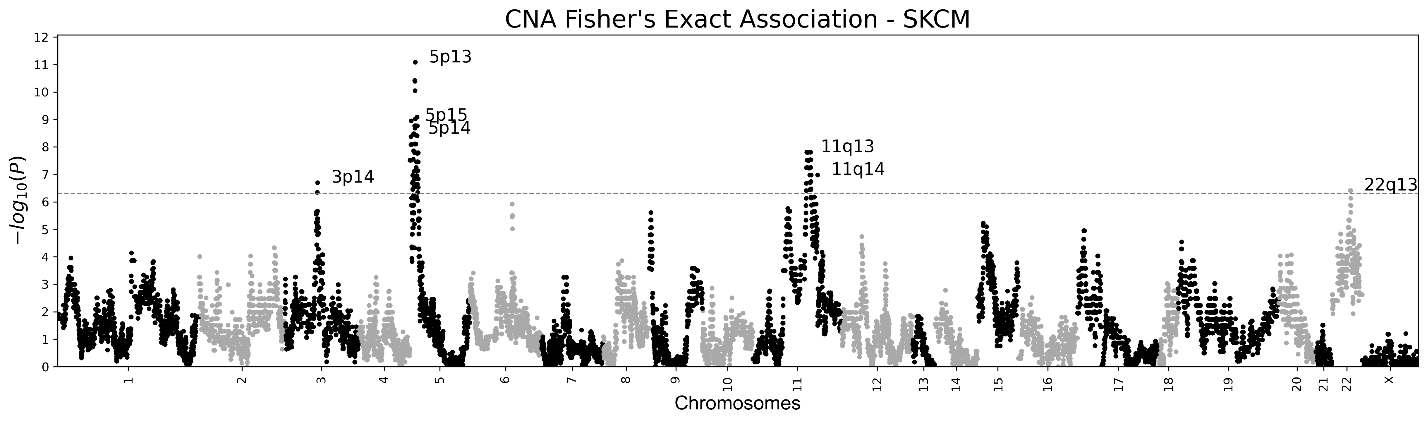

**b**
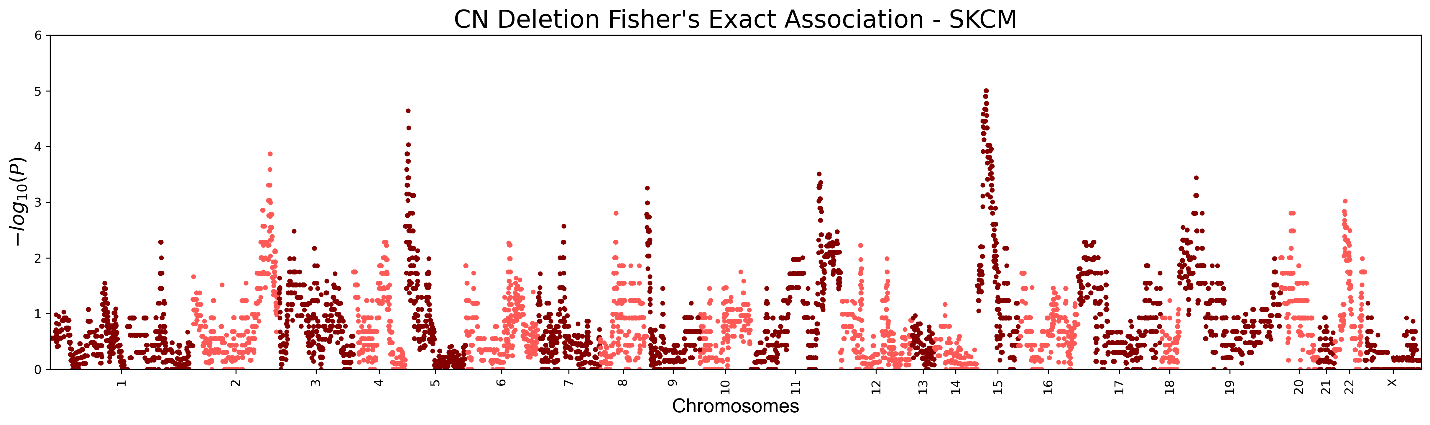

**c**
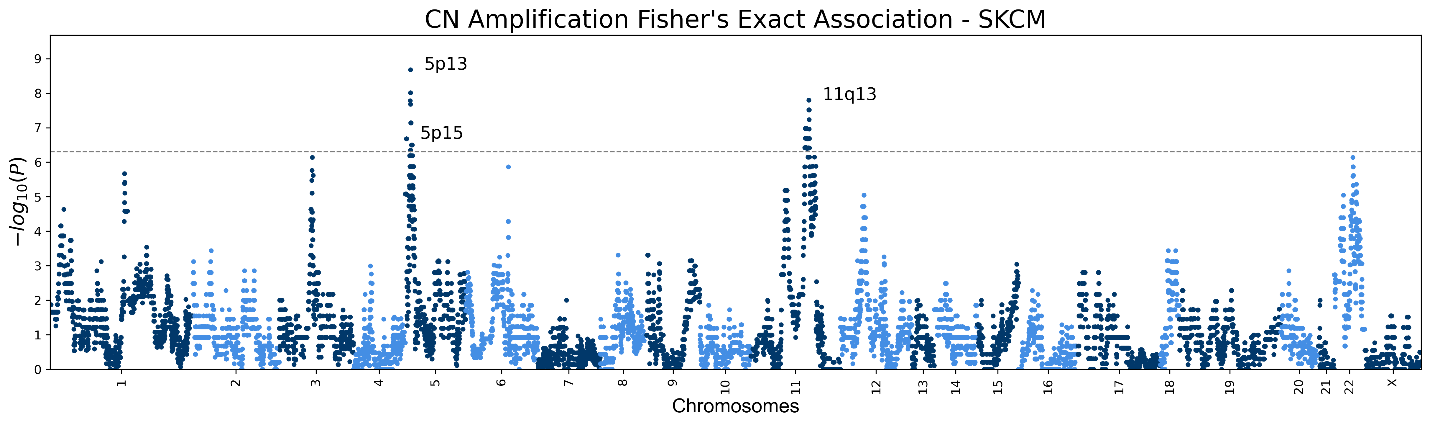

**Figure S19. STAD Manhattan plots**

Genic Manhattan plot over Fisher’s exact test p-values between STAD chromoanagenesis samples and non-chromoanagenesis samples. **(a)** Manhattan CNA (deletion or amplification) plot. **(b)** Manhattan Deletion plot. **(c)** Manhattan Amplification plot.

**a**
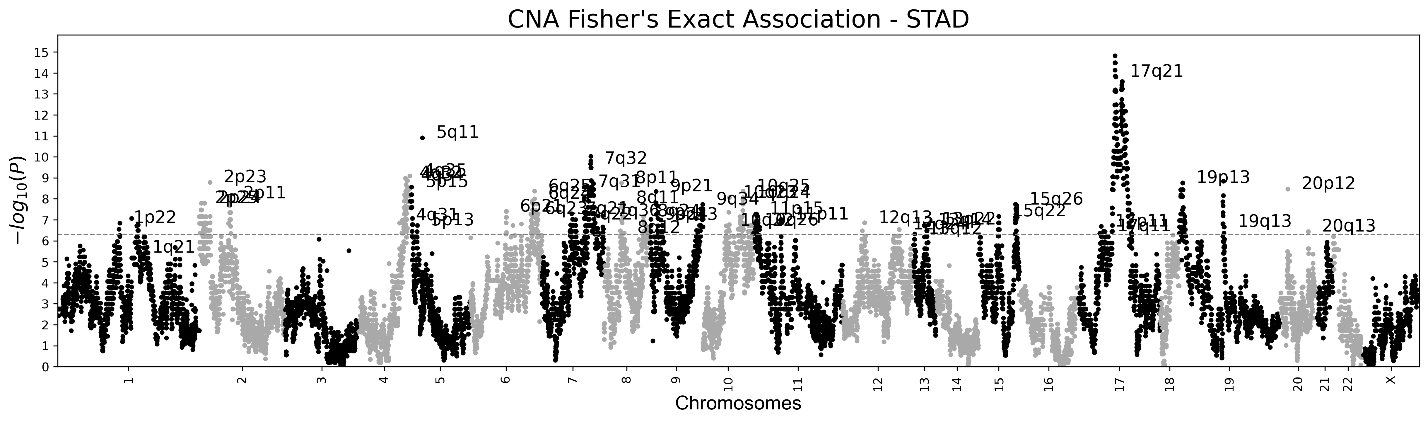

**b**
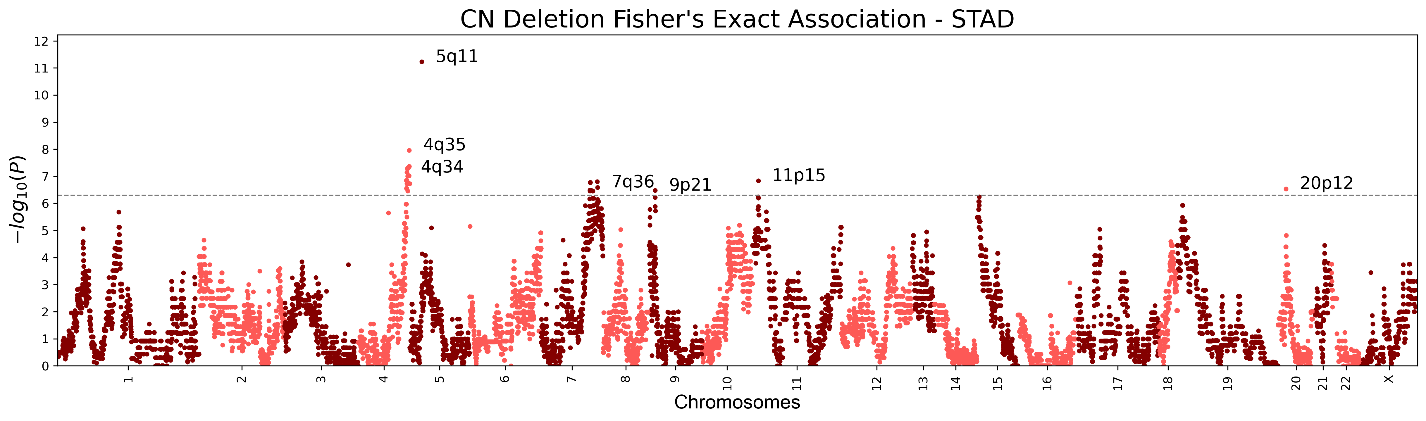

**c**
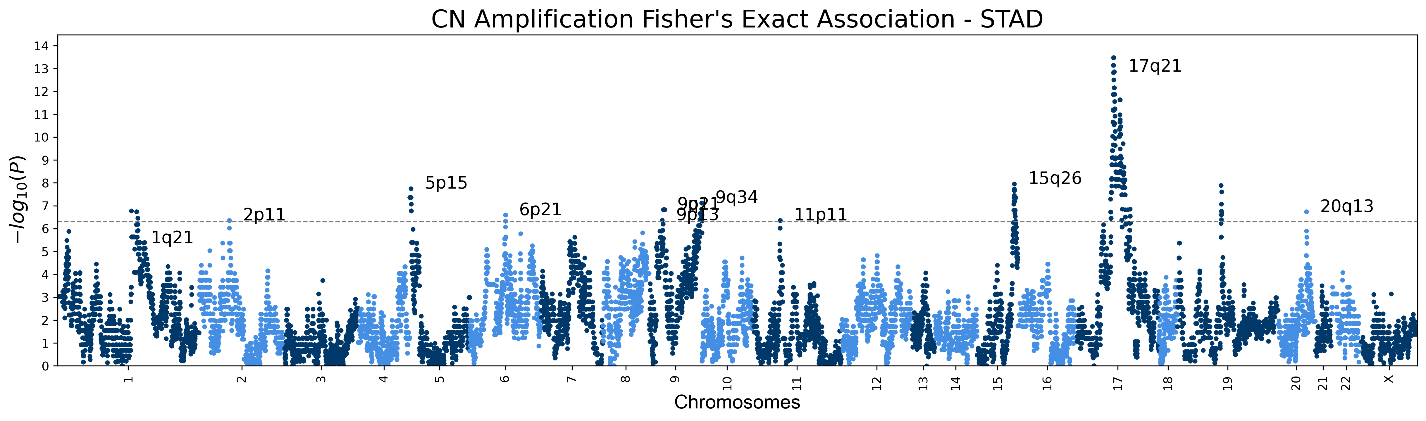

**Figure S20. UCEC Manhattan plots**

Genic Manhattan plot over Fisher’s exact test p-values between UCEC chromoanagenesis samples and non-chromoanagenesis samples. **(a)** Manhattan CNA (deletion or amplification) plot. **(b)** Manhattan Deletion plot. **(c)** Manhattan Amplification plot.

**a**
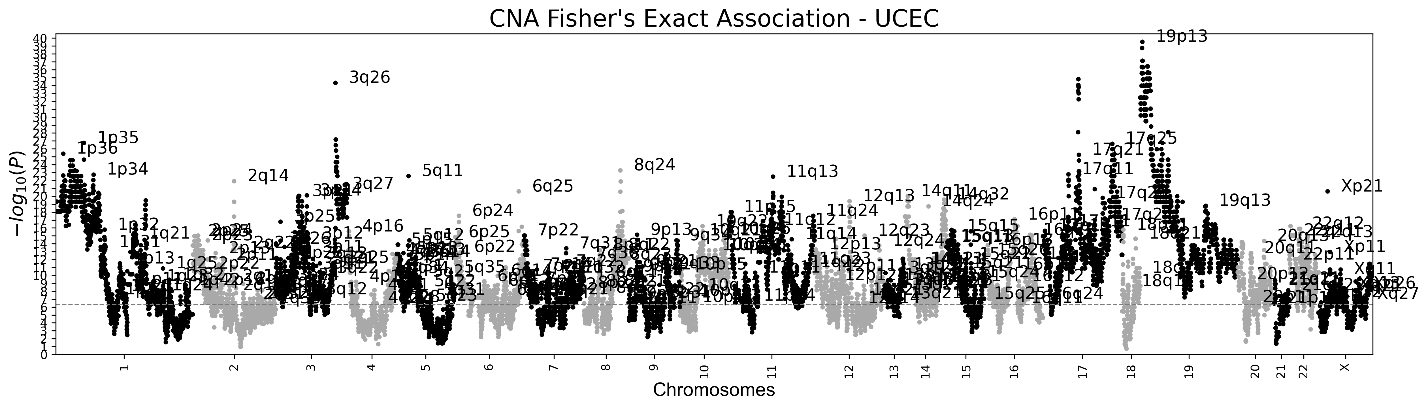

**b**
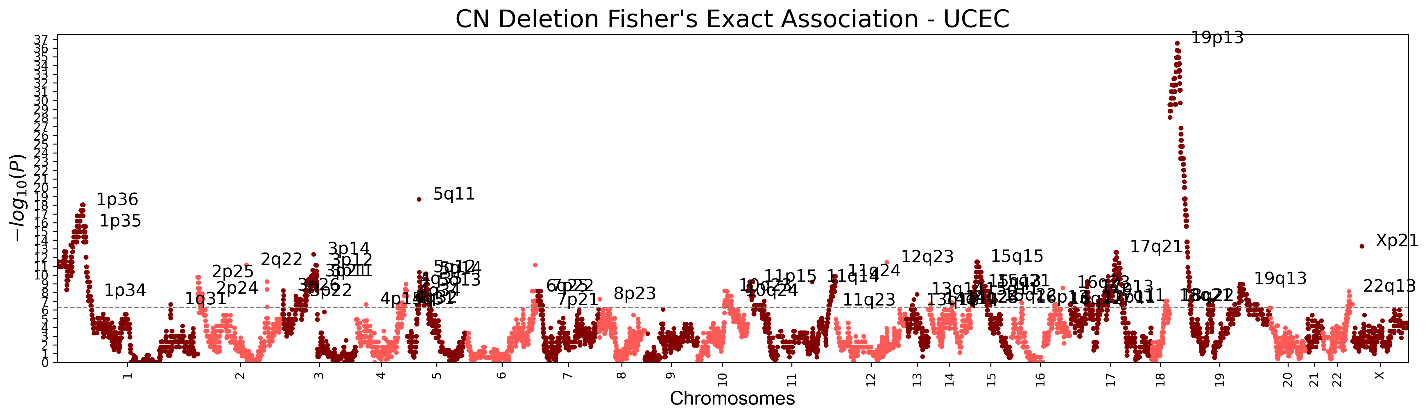

**c**
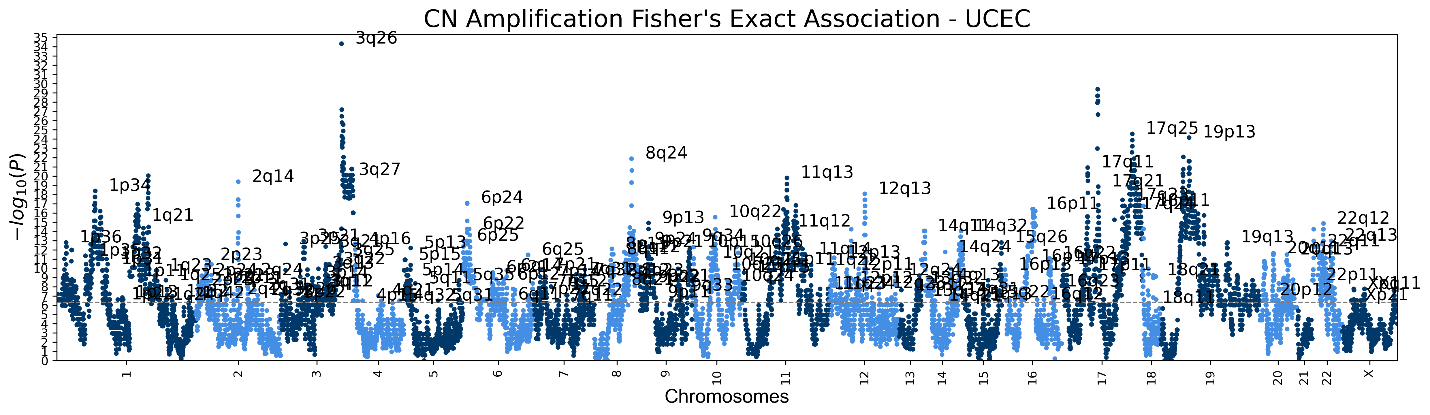

**Figure S21.**  **(a)** Kaplan-Meier Overall survival rate estimate for chromoanagenesis in BLCA. **(b)** Cox regression hazard ratio, for chromoanagenesis, age and gender in BLCA patients.

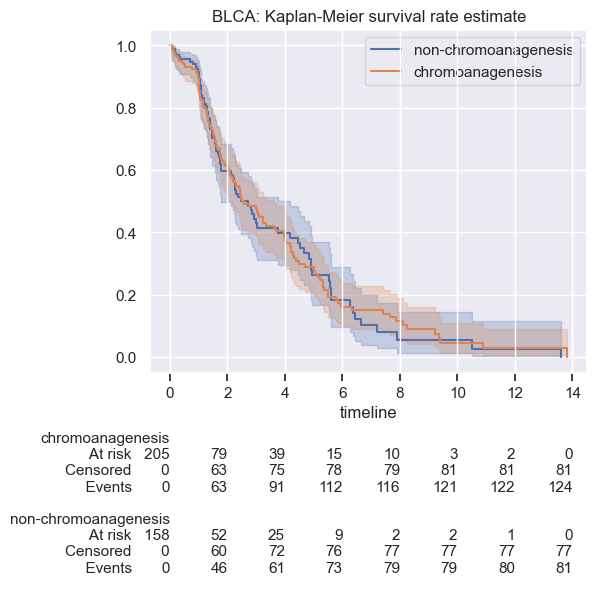
**a**

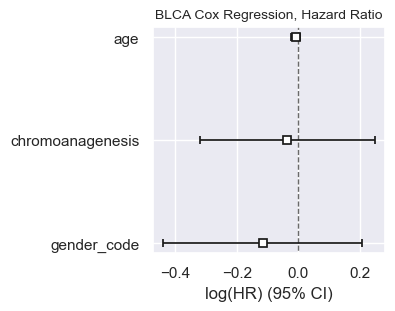
**b**

**Figure S22.**  **(a)** Kaplan-Meier Overall survival rate estimate for chromoanagenesis in BRCA. **(b)** Cox regression hazard ratio, for chromoanagenesis, age and gender in BRCA patients.

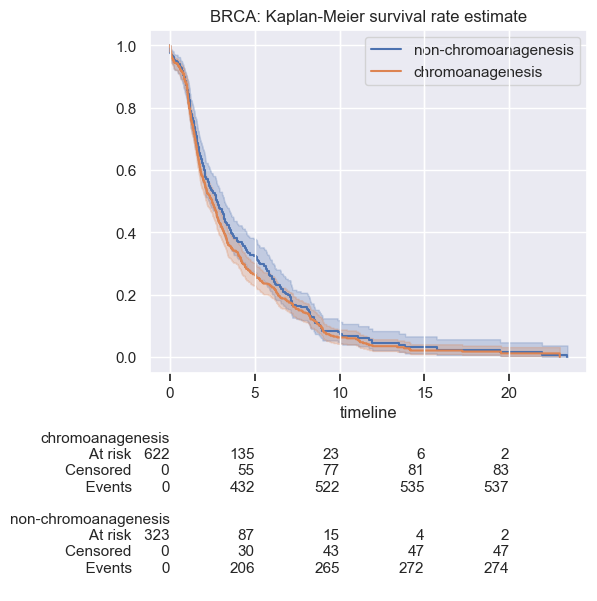

**a**

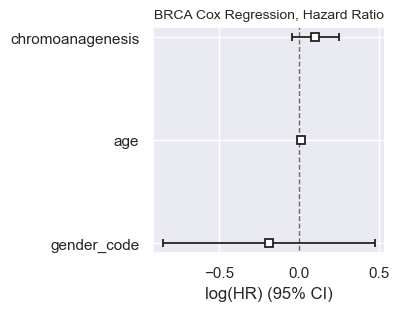
**b**

**Figure S23.**  Kaplan-Meier Overall survival rate estimate for chromoanagenesis in CESC.

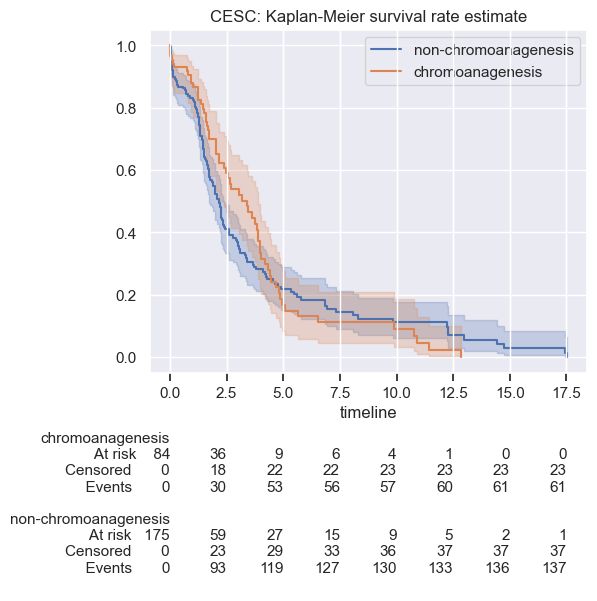

**Figure S24.**  **(a)** Kaplan-Meier Overall survival rate estimate for chromoanagenesis in COAD. **(b)** Cox regression hazard ratio, for chromoanagenesis, age and gender in COAD patients.

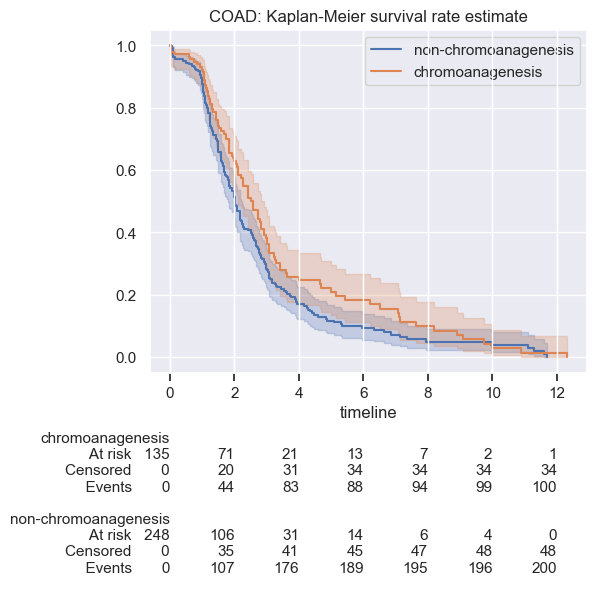
**a**

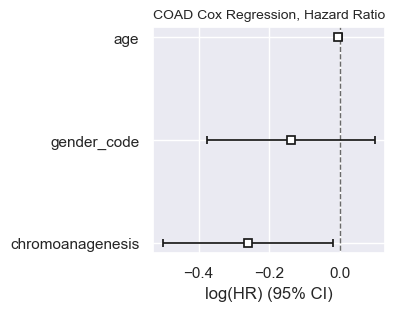
**b**

**Figure S25.**  **(a)** Kaplan-Meier Overall survival rate estimate for chromoanagenesis in ESCA. **(b)** Cox regression hazard ratio, for chromoanagenesis, age and gender in ESCA patients.

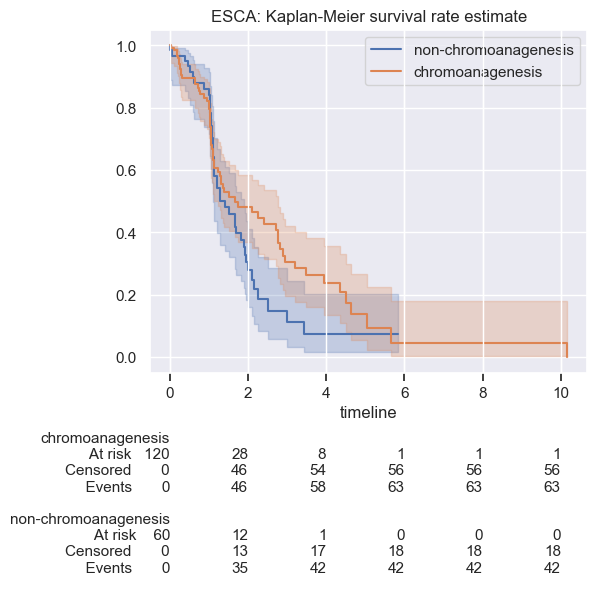

**a**

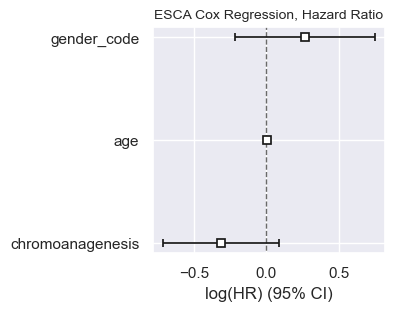

**b**

**Figure S26.**  **(a)** Kaplan-Meier Overall survival rate estimate for chromoanagenesis in GBM. **(b)** Cox regression hazard ratio, for chromoanagenesis, age and gender in GBM patients.

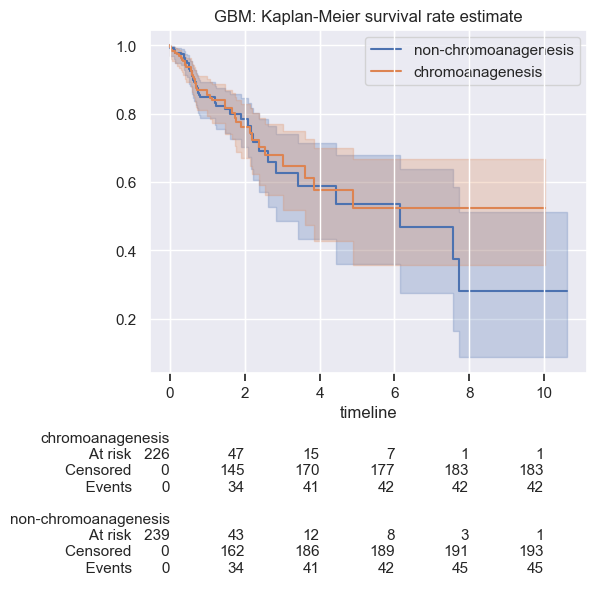
**a**

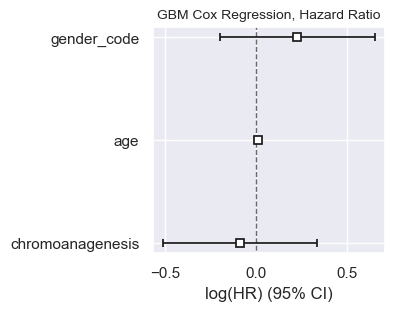

**b**

**Figure S27.**  **(a)** Kaplan-Meier Overall survival rate estimate for chromoanagenesis in HNSC. **(b)** Cox regression hazard ratio, for chromoanagenesis, age and gender in HNSC patients.

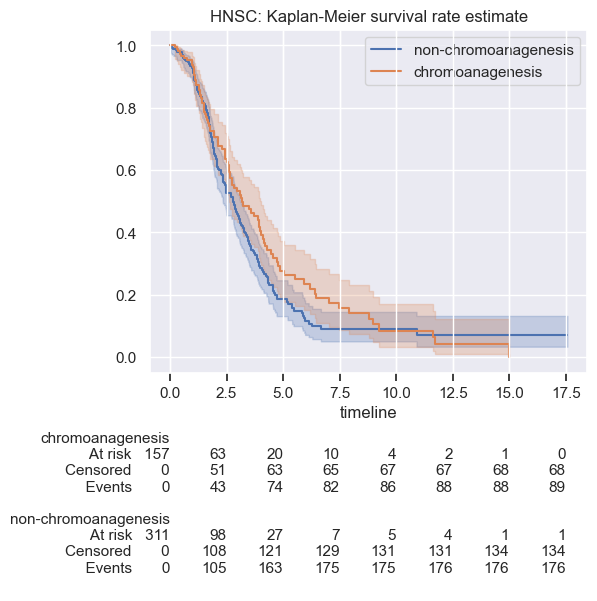
**a**

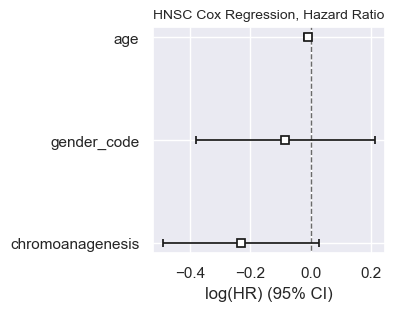

**b**

**Figure S28.**  **(a)** Kaplan-Meier Overall survival rate estimate for chromoanagenesis in KIRC. **(b)** Cox regression hazard ratio, for chromoanagenesis, age and gender in KIRC patients.

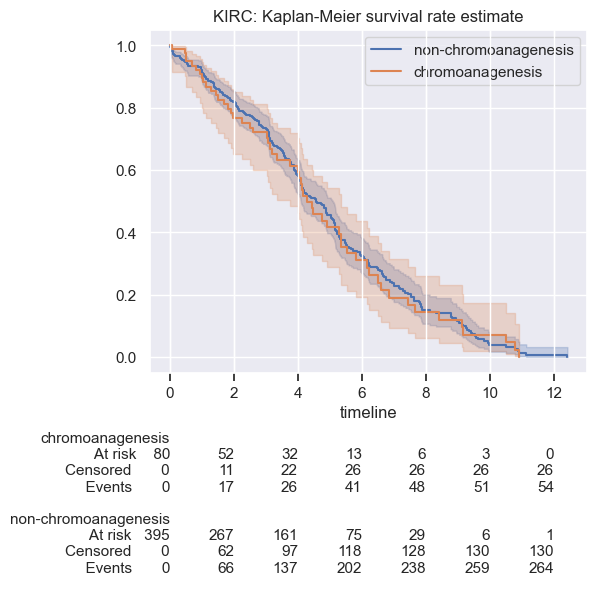

**a**

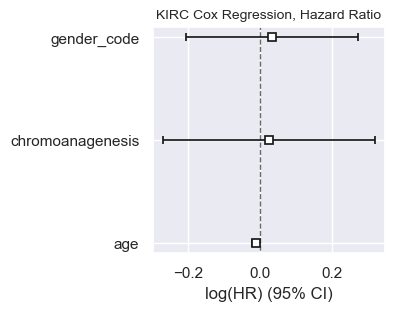

**b**

**Figure S29.**  **(a)** Kaplan-Meier Overall survival rate estimate for chromoanagenesis in LGG. **(b)** Cox regression hazard ratio, for chromoanagenesis, age and gender in LGG patients.

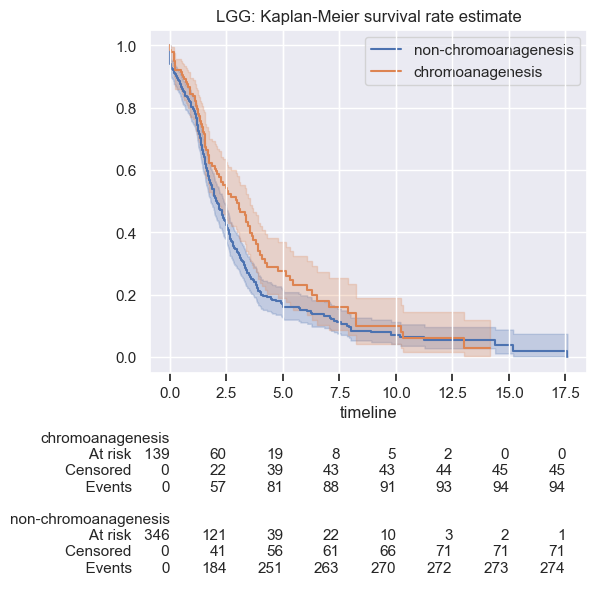
**a**

**b**
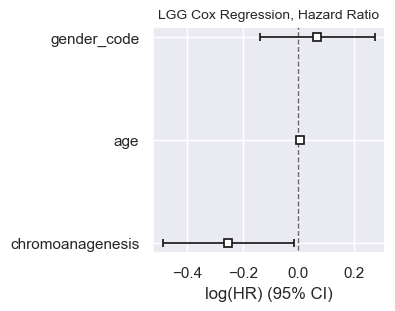

**Figure S30.**  **(a)** Kaplan-Meier Overall survival rate estimate for chromoanagenesis in LIHC. **(b)** Cox regression hazard ratio, for chromoanagenesis, age and gender in LIHC patients.

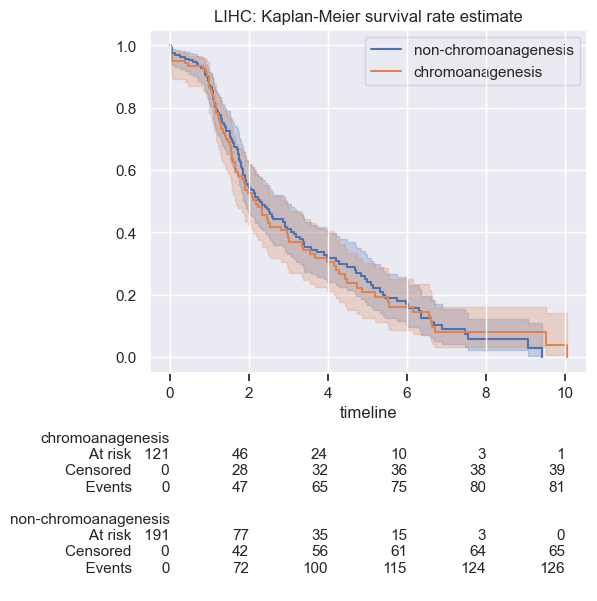
**a**

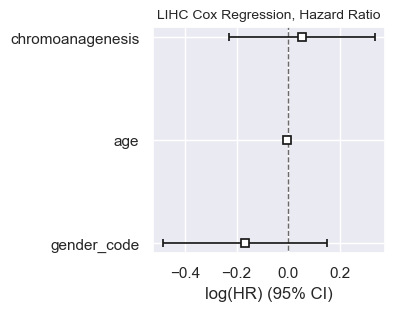

**b**

**Figure S31.**  **(a)** Kaplan-Meier Overall survival rate estimate for chromoanagenesis in LUAD. **(b)** Cox regression hazard ratio, for chromoanagenesis, age and gender in LUAD patients.

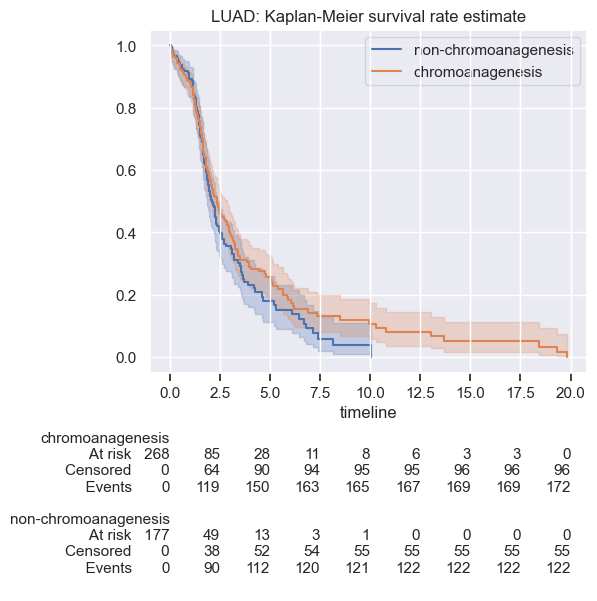

**a**

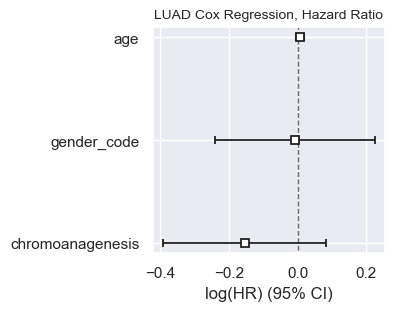

**b**

**Figure S32.**  **(a)** Kaplan-Meier Overall survival rate estimate for chromoanagenesis in LUSC. **(b)** Cox regression hazard ratio, for chromoanagenesis, age and gender in LUSC patients.

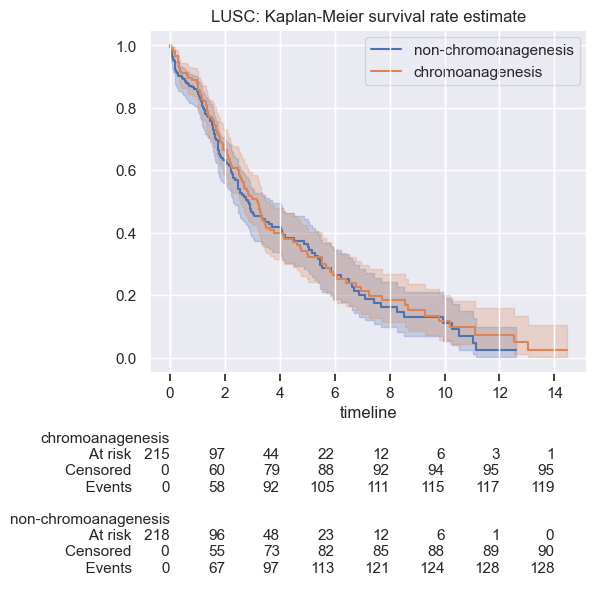
**a**

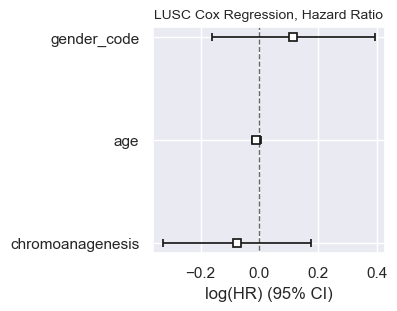
**b**

**Figure S33.**  Kaplan-Meier Overall survival rate estimate for chromoanagenesis in OV.

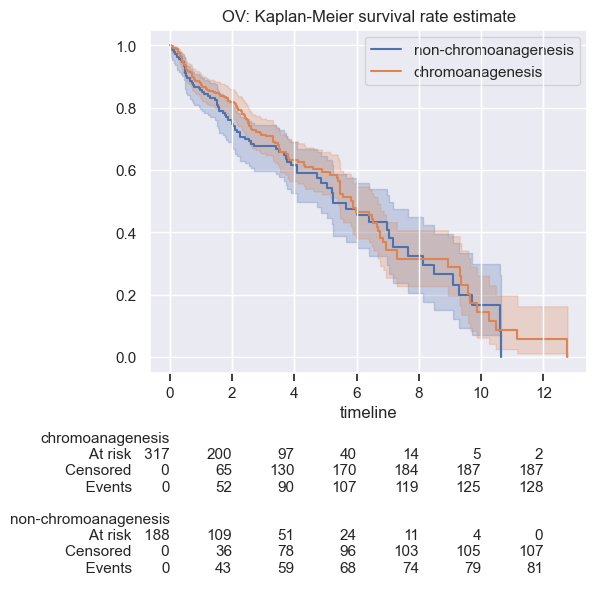

**Figure S34.**  **(a)** Kaplan-Meier Overall survival rate estimate for chromoanagenesis in PAAD. **(b)** Cox regression hazard ratio, for chromoanagenesis, age and gender in PAAD patients.

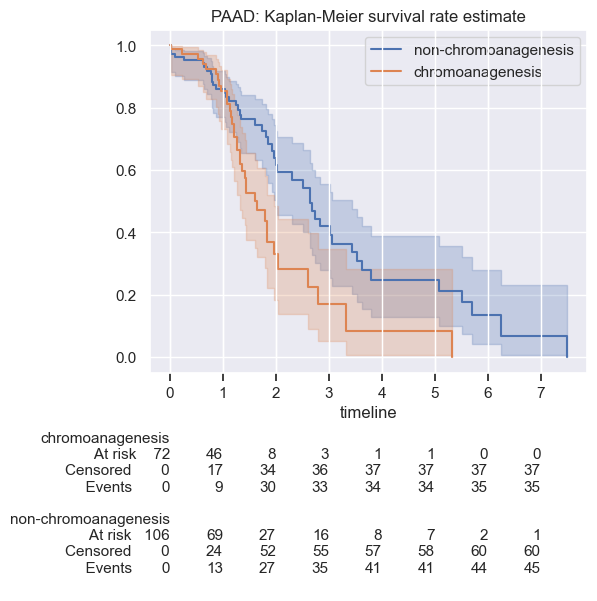
**a**

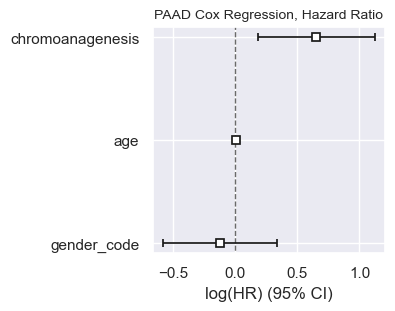

**b**

**Figure S35.** Kaplan-Meier Overall survival rate estimate for chromoanagenesis in PRAD.

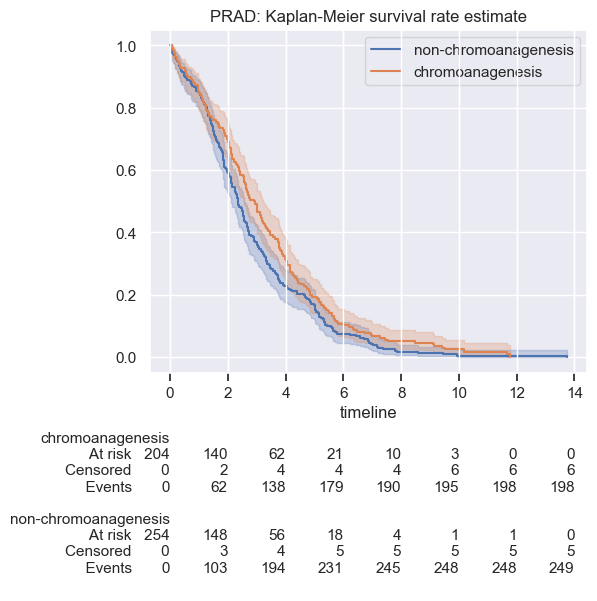

**Figure S36.**  **(a)** Kaplan-Meier Overall survival rate estimate for chromoanagenesis in READ. **(b)** Cox regression hazard ratio, for chromoanagenesis, age and gender in READ patients.

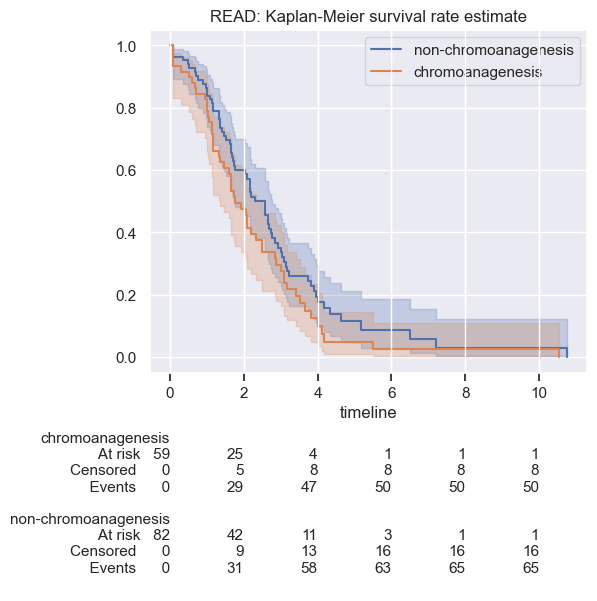
**a**

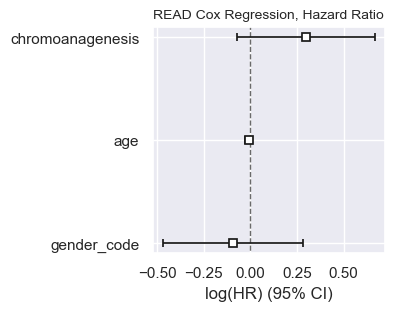
**b**

**Figure S37.**  **(a)** Kaplan-Meier Overall survival rate estimate for chromoanagenesis in SARC. **(b)** Cox regression hazard ratio, for chromoanagenesis, age and gender in SARC patients.

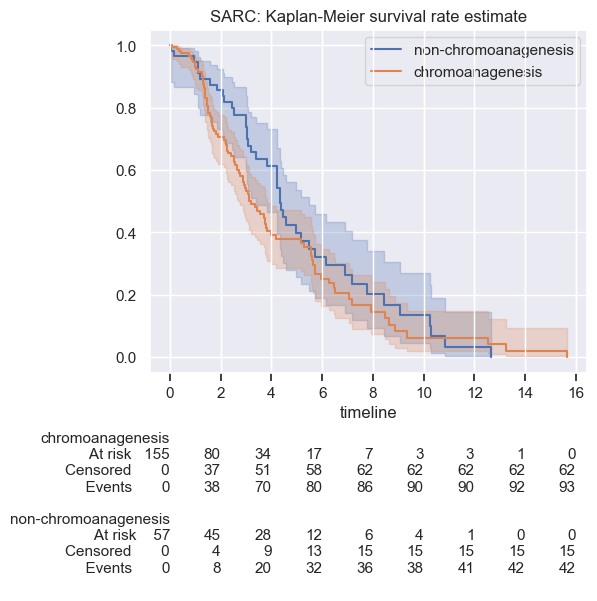
**a**

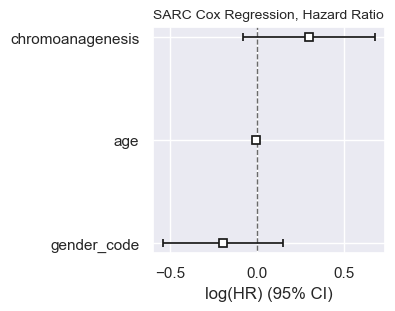
**b**

**Figure S38. (a)** Kaplan-Meier Overall survival rate estimate for chromoanagenesis in SKCM. **(b)** Cox regression hazard ratio, for chromoanagenesis, age and gender in SKCM patients.

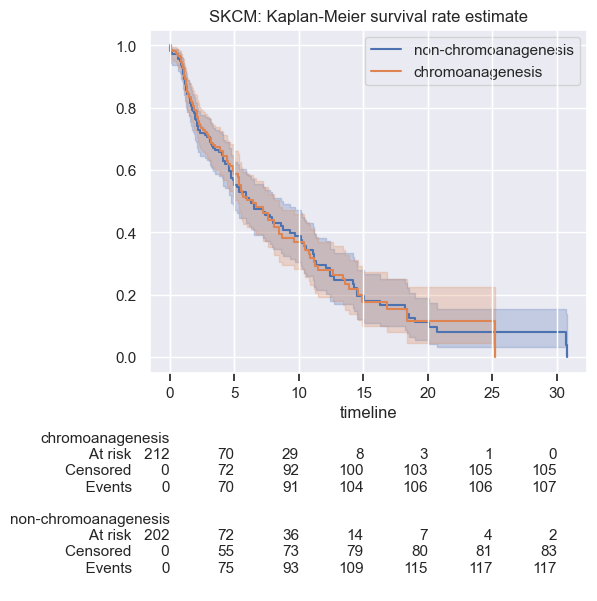

**a**

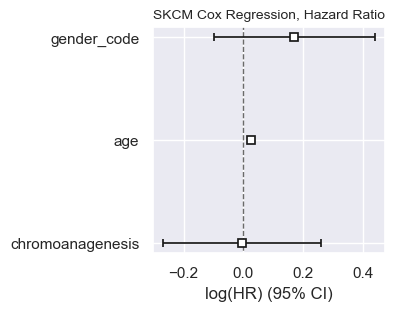

**b**

**Figure S39.**  **(a)** Kaplan-Meier Overall survival rate estimate for chromoanagenesis in STAD. **(b)** Cox regression hazard ratio, for chromoanagenesis, age and gender in STAD patients.

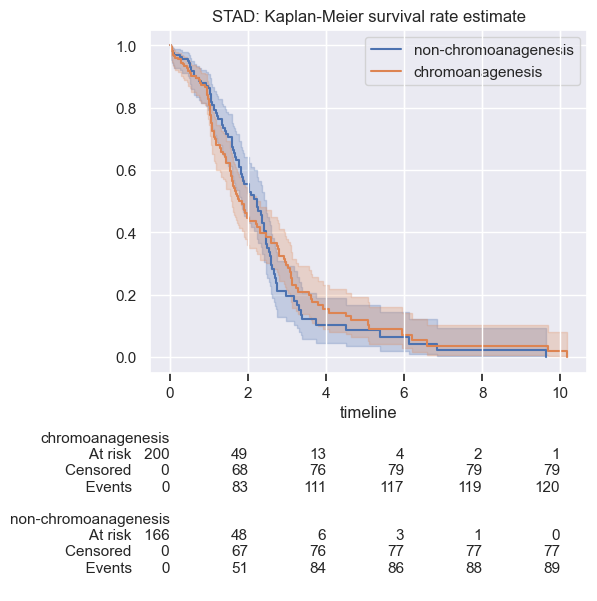

**a**

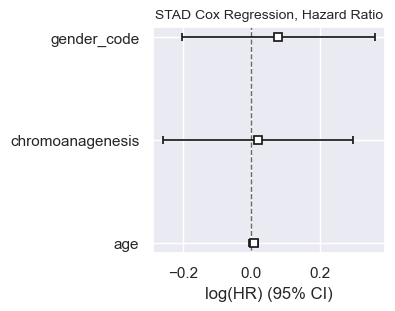

**b**

**Figure S40.** Kaplan-Meier Overall survival rate estimate for chromoanagenesis in UCEC.

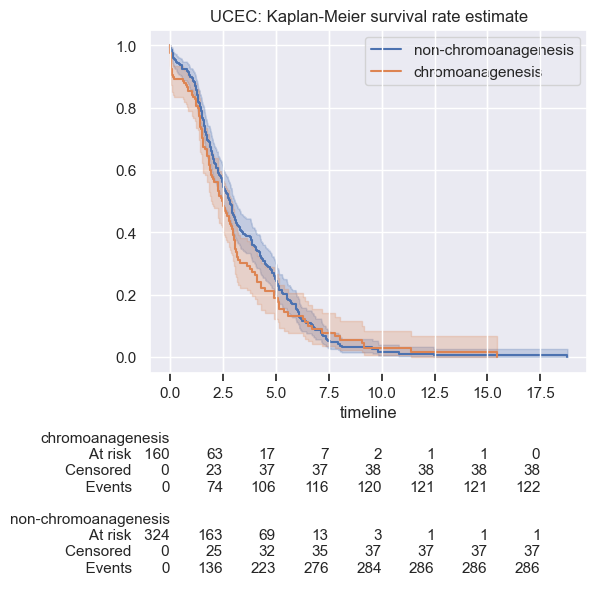
